# A Theory of Synaptic Transmission

**DOI:** 10.1101/2020.12.17.423159

**Authors:** Bin Wang, Olga K. Dudko

## Abstract

Rapid and precise neuronal communication is enabled through a highly synchronous release of signaling molecules neurotransmitters into the synaptic cleft within just milliseconds of the action potential. Yet neurotransmitter release lacks a theoretical framework that is both phenomenologically accurate and mechanistically realistic. Here, we formulate an analytic theory of the action-potential-triggered neurotransmitter release at a chemical synapse. The theory captures general principles of synaptic transmission while generating concrete predictions for particular synapses. A universal scaling in synaptic transmission is established, and demonstrated through a collapse of experimental data from different synapses onto a universal curve. The theory shows how key characteristics of synaptic function – plasticity, fidelity, and efficacy – emerge from molecular mechanisms of neurotransmitter release machinery.

## Introduction

Neurons communicate across special junctions – synapses – using neurotransmitter molecules as a chemical signal (***Südhof, 2013***). Release of neurotransmitters into the synaptic gap occurs when neurotransmitter-loaded vesicles fuse with the membrane of the presynaptic (transmitting) neuron in response to calcium influx during an action potential “spike”. Synaptic vesicle fusion is remarkably fast and precise: both the duration of fusion and the time between the trigger and fusion initiation are less than a millisecond (***Katz and Miledi, 1965***).

The electrical propagation of information along the axon of the presynaptic neuron (the pre-transmission stage) and the response of the postsynaptic neuron to the chemical signal (the post-transmission stage) have been described by theories that capture phenomenology while connecting to microscopic mechanisms (***Hodgkin and Huxley, 1952; Destexhe et al., 1994***). However, neurotransmitter release, which enables the synaptic transmission itself, lacks a theory that is both phenomenologically accurate and microscopically realistic (***Stevens, 2000***). This void contrasts with detailed experiments, which have revealed the molecular constituents involved. The key to speed and precision of neurotransmitter release is a calcium-triggered conformational transition in SNAREs (<u>s</u>oluble <u>N</u>-ethylmaleimide sensitive factor <u>a</u>ttachment protein <u>re</u>ceptors) (***Baker and Hughson, 2016; Brunger et al., 2018; Kaeser and Regehr, 2014***). The free energy released during the conformational transition is harnessed by SNAREs to pull the membranes of the vesicle and the cell together, reducing the high kinetic barriers that otherwise hinder fusion. Fusion culminates in the release of neurotransmitters from vesicles into the synaptic cleft (Fig. 1A).

**Figure 1.**
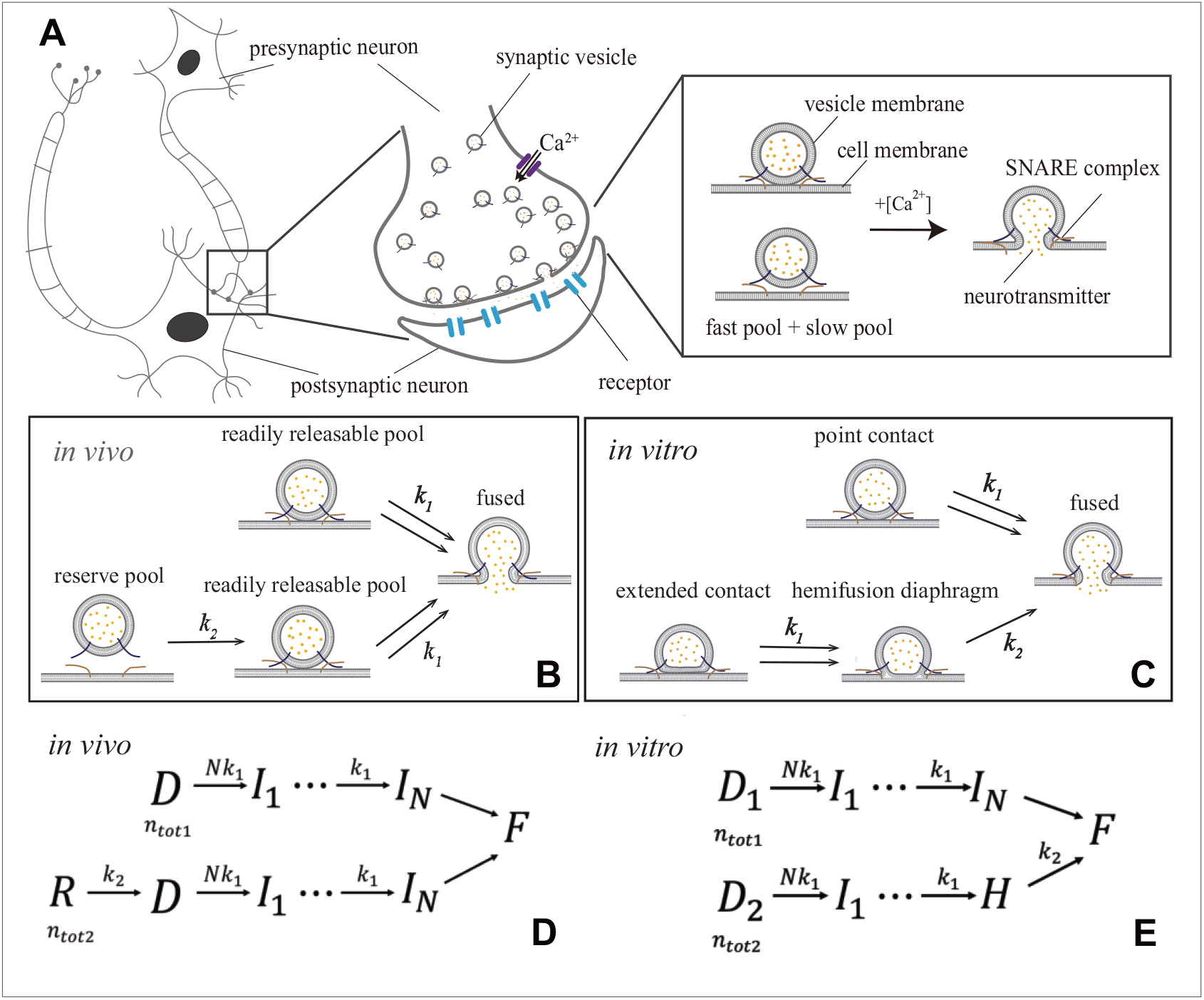
Synaptic transmission *in vivo* and *in vitro*. (A) Release of neurotransmitters into the synaptic cleft (diameter ~ 1 − 20*μm*) occurs when neurotransmitter-loaded vesicles (diameter ~ 30*nm*) fuse with the presynaptic cell membrane in response to *Ca*^2+^ influx during an action potential. Fusion is facilitated by SNARE protein complexes and proceeds via two parallel pathways that originate in the “fast” and “slow” vesicle pools. (B and C) Fusion stages *in vivo* and *in vitro*. SNARE conformational transition constitutes the fast step, *k*_1_. Vesicle transfer from the reserve pool to the readily releasable pool (RRP) *in vivo* and escape from the hemifusion diaphragm *in vitro* constitute the slow step, *k*_2_. (D and E) Reaction schemes for (B) and (C). *In vivo*, state *R* represents the reserve pool, *D* the RRP, *I*_*i*_ the state with *i* SNAREs that underwent conformational transitions, *F* the fused state. *In vitro*, *D*_1_ and *D*_2_ represent docked vesicles with point- and extended-contact morphologies, *H* the hemifusion diaphragm. Mathematical equivalence of the reaction schemes *in vivo* and *in vitro* enables the treatment through a unifying theory.

Here, we present a theory of synaptic transmission, which quantitatively reproduces a wide range of data from fluorescence experiments *in vitro* (***Kyoung et al., 2011; Diao et al., 2012***) and electrophysiological experiments *in vivo* (***Schneggenburger and Neher, 2000; Lou et al., 2005; Bollmann et al., 2000; Wölfel et al., 2007; Kochubey et al., 2009; Duncan et al., 2010; Miki et al., 2018***). The theory yields analytic expressions for measurable quantities, which enables a direct fit to the data. The fit extracts parameters of synaptic fusion machinery: activation barriers and rates of SNARE conformational transitions at any calcium concentration, the size of vesicle pools, and the number of SNAREs necessary for fusion. The derived expression for the calcium-dependent release rate quantifies the remarkable temporal precision of neurotransmitter release. A universal relation for the peak release rate is established, and demonstrated through a collapse of data from a variety of synapses onto a universal curve. The theory is further applied to relate the properties of neurotransmitter release machinery to short-term plasticity, rate/fidelity trade-off, and synaptic efficacy, thus providing a mapping from molecular constituents to functions in synaptic transmission.

## Results

### Theory

We start from the observation that published data on neurotransmitter release for different synapses and experimental setups (***Kyoung et al., 2011; Diao et al., 2012; Schneggenburger and Neher, 2000; Lou et al., 2005; Bollmann et al., 2000; Miki et al., 2018; Duncan et al., 2010; Kochubey et al., 2009; Wölfel et al., 2007***) can all be encompassed by a unifying kinetic scheme:

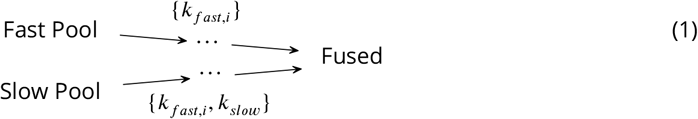

In this kinetic scheme, synaptic vesicle fusion proceeds through two parallel reaction pathways. Both pathways contain fast steps of rate constants {*k*_*fast,i*_}. One of the pathways contains an additional, slow, step of rate constant *k*_*slow*_ ≪ {*k*_*fast,i*_}. The pathways originate in the “fast” and “slow” vesicle pools of sizes *n*_*tot*1_ and *n*_*tot*2_, respectively (Eq. (1)). The interpretations of the fast and slow steps as well as the individual states in this unifying kinetic scheme for different experimental setups are detailed below.

In the context of *in vivo* experiments (***Schneggenburger and Neher, 2000; Lou et al., 2005; Bollmann et al., 2000; Miki et al., 2018; Duncan et al., 2010; Kochubey et al., 2009; Wölfel et al., 2007***), Eq. (1) concretizes into the kinetic scheme in Fig. 1B and D. The fast pool represents the readily releasable pool (RRP) comprised of vesicles that are docked on the presynaptic terminal (state *D*) and fuse readily upon *Ca*^2+^ influx (***Kaeser and Regehr, 2017***). The slow pool represents the reserve pool (state *R*), which supplies vesicles to the RRP (*R* → *D*) with slow rate *k*_2_. Fusion of an RRP vesicle (…→ *F*) requires *N* SNAREs tethering the vesicle at the cell membrane to concurrently undergo a conformational transition. This transition is *Ca*^2+^-dependent and involves a single rate-limiting step (***Hui et al., 2005***) of rate constant *k*_1_([*Ca*^2+^]).

In the context of *in vitro* experiments (***Kyoung et al., 2011; Diao et al., 2012***), Eq. (1) becomes the kinetic scheme in Fig. 1C and E. All vesicles are initially docked (states *D*_1_ and *D*_2_) but adopt different morphologies (Fig. 1C) and, consequently, fuse through different pathways (***Gipson et al., 2017***). Vesicles in a point contact with the membrane (state *D*_1_) fuse rapidly upon *Ca*^2+^-triggered SNARE conformational transition, mimicking RRP vesicles *in vivo*. Vesicles in an extended contact (state *D*_2_) become trapped in a hemifusion diaphragm intermediate (state *H*), escape from which (*H* → *F*) constitutes the slow step *k*_2_.

In all these experiments, the delay due to steps *I*_*N*_ → *F* is negligible compared to both fast and slow steps *k*_1_ and *k*_2_. Note that a scheme with *N* concurrent steps of rates *k*_1_ (Fig. 1B and C) is equivalent to a scheme with *N* sequential steps of rates *Nk*_1_, (*N* − 1)*k*_1_,…,*k*_1_ (Fig. 1D and E).

Despite the differences in fusion stages *in vivo* and *in vitro* described above, mathematical equivalence of the corresponding kinetic schemes enables their treatment through a unifying theory. We will assume that the calcium influx is triggered by an action potential that arrives at the presynaptic terminal at *t* = 0. The microsecond timescales (much faster than neurotransmitter release) of the opening of voltage-gated *Ca*^2+^ channels and diffusion of *Ca*^2+^ ions across the active zone justify treating the [*Ca*^2+^] rising as instantaneous. Since the typical width of [*Ca*^2+^] profile is ~ 1 − 10ms (***Bean, 2007***) while most vesicles fuse within *t* ~ 100*μs* (***Katz and Miledi, 1965***), [*Ca*^2+^] can be treated as approximately constant during the fusion process. The theory is thus applicable both for step-like and for spike-like [*Ca*^2+^] profiles, as well as for spike trains if the recording time is shorter than the timescale 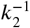 of RRP replenishment. With the above assumptions, the theory is developed in detail in Appendix 1. Below we present analytic expressions derived from the theory for the key quantities that are measured in the experiments that probe synaptic transmission at the single-synapse level *in vivo* and *in vitro*. These expressions relate experimentally measurable characteristics of synaptic transmission to the microscopic parameters of synaptic release machinery and thus enable the extraction of these parameters through a fit to experimental data.

An informative characteristic of synaptic transmission is the average release rate. Defined as the average (over an ensemble of repeated stimuli) rate of change in the number of fused vesicles, this quantity “encodes” information about the kinetic signatures and is the most commonly reported characteristic in experiments on the kinetics of neurotransmitter release (***Schneggenburger and Neher, 2000; Bollmann et al., 2000; Kyoung et al., 2011; Diao et al., 2012***). The rate equations for the kinetic scheme in Eq. (1) yield the exact solution for the average release rate:

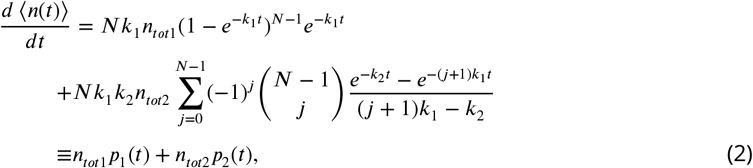

where *p*_1,2_(*t*) are the probability distributions for the fusion time in the fast and slow pathways, *N* is the necessary number of SNAREs, and *n*_*tot*1_ and *n*_*tot*2_ are the sizes of the fast and slow pools, respectively. We use the standard notation for binomial coefficient 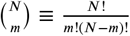.

In practice, the average release rate is obtained from the average cumulative release ⟨*n*(*t*)⟩, which is defined as the average number of vesicles fused by time *t* and can be measured directly through fluorescence imaging (***Kyoung et al., 2011; Diao et al., 2012***) or electrophysiological recording (***Schneggenburger and Neher, 2000; Lou et al., 2005; Bollmann et al., 2000; Wölfel et al., 2007; Kochubey et al., 2009; Duncan et al., 2010; Miki et al., 2018***) on the postsynaptic neuron. Integrating Eq. (2) yields the exact solution for average cumulative release:

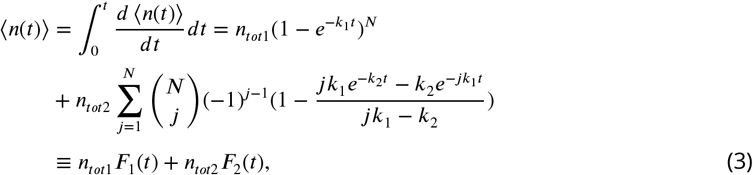

where 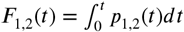 are cumulative distributions for the fusion time in the fast and slow pathways. *In vivo*, *F*_1_(*t* = *T*) is the fusion probability for an RRP vesicle after an action potential of duration *T* (***Neher, 2015; Miki et al., 2018; Malagon et al., 2016***). We also derive the full probability distribution of cumulative release (Appendix 1), which, although at present is challenging to measure experimentally, contains more information than the average values (Eqs. (2)–(3)).

Previous experiments indicate a separation of timescales, *k*_2_ ≪ *k*_1_ (***Neher, 2010; Kaeser and Regehr, 2014***), which yields useful asymptotic behaviors. At short times, *t ≪* 1/*k*_1_, 1/*k*_2_, the release rate in Eq. (2) is 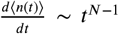, which can be readily fit to data to extract the number of SNAREs *dt* necessary for fusion. At intermediate times, 1/*k*_1_ ≪ *t* ≪ 1/*k*_2_, cumulative release in Eq. (3) becomes ⟨*n*(*t*)⟩ ≈ *n*_*tot*1_ + *n*_*tot*2_*k*_2_*t*, which can be used to determine the RRP size, *n*_*tot*1_, by extrapolation (***Neher, 2015***). At long times, *t* ~ 1/*k*_2_ ≫ 1/*k*_1_, we get 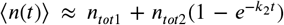. As expected, the cumulative release on the intermediate and long timescales is independent of the number *N* and conformational rate *k*_1_ of SNAREs as all the fast steps have been completed.

A measure of sensitivity of a synapse to [*Ca*^2+^] is the peak release rate (***Schneggenburger and Neher, 2000; Lou et al., 2005; Bollmann et al., 2000***). The time at which the peak is reached is found from Eq. (2) using 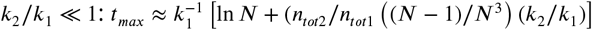. The peak release rate is then

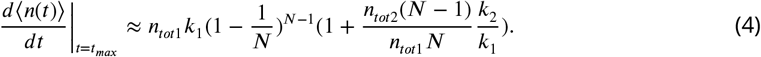

Applicability of the framework in Eqs. (2)–(4) requires an explicit form for the calcium-dependence of the rate constant of SNARE conformational transition, *k*_1_([*Ca*^2+^]). We utilize Kramers formalism of reaction kinetics (***Kramers, 1940; Dudko et al., 2006***), which treats a conformational transition as thermal escape over a free energy barrier along a reaction coordinate *Q*. In the present context, the role of *Q* is fulfilled by the average number of *Ca*^2+^ ions bound to a SNARE at a given [*Ca*^2+^], assuming that this average follows the dynamics of SNARE conformational degree of freedom. The generic shape of the free energy profile with a barrier that separates the two conformational states of a SNARE complex is captured by a cubic polynomial. The effect of calcium on the free energy profile is incorporated analogous to the *pH*-dependence of Gibbs free energy of a protein (***Schaefer et al., 1997; Zhang and Dudko, 2015***). The Kramers rate is then

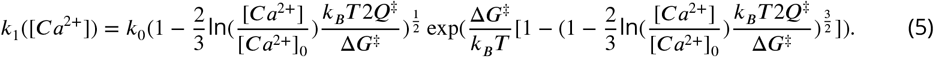

Here, *k*_0_, Δ*G*^‡^ and *Q*^‡^ are the rate constant and the height and width of the activation barrier for SNARE conformational transition at a reference calcium concentration [*Ca*^2+^]_0_. The argument of exp(…) is the change in the barrier height at a given [*Ca*^2+^]. Since the release rate (Eq. (2)) and its peak (Eq. (4)) are proportional to *k*_1_([*Ca*^2+^]), both quantities are exponentially sensitive to [*Ca*^2+^] (Appendix 1 Fig. 1A). This exponentially strong sensitivity of the release rate to calcium concentration explains the precisely timed character of synaptic release: synaptic fusion machinery turns on rapidly upon *Ca*^2+^ influx during the action potential and terminates rapidly upon *Ca*^2+^ depletion.

Equations (2) and (5) reveal that *N* = 2 SNAREs per vesicle provide the optimal balance between stability and temporal precision of release dynamics (***Sinha et al., 2011***). Indeed, at *N* = 1, the release is hypersensitive to sub-millisecond [*Ca*^2+^] fluctuations caused by stochastic opening of *Ca*^2+^ channels (note the high release rate on sub-millisecond timescale at *N* = 1 in Appendix 1 – Fig. 1B). On the other hand, *N* > 2 delay the peak of release following an action potential. The optimality of *N* = 2 is further supported by the least squares fit of the experimental data (***Kochubey et al., 2009***) to Eq. (3) with different values of *N*: *N* = 2 results in the smallest fitting errors for all calcium concentrations used in the experiment (Appendix 3). Given the limited experimental data available, below we fix *N* at the optimal value of 2 in order to constrain our fits. However, provided sufficient data is available for the fit, *N* in the present theory can serve as a free parameter. For *N* = 2, Eq. (4) for the peak release rate simplifies to

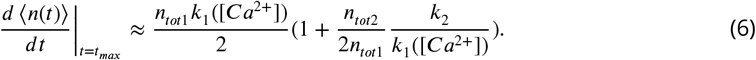

The analytic results, Eqs. (4) and (5), allow us to establish a universal relation for the sensitivity of a synapse to the strength of the trigger:

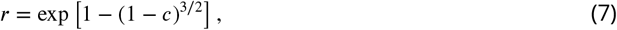

where 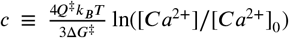 and 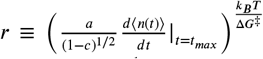 are the dimensionless calcium concentration and peak release rate, and 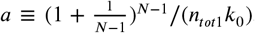. If the scaling law in Eq.(7) indeed captures universal principles of synaptic transmission, data from different synapses should collapse onto the curve given by Eq. (7). This prediction is tested below.

A postsynaptic response to the action potential events is measured by the peak value of the postsynaptic current (PSC). Using the well-established conductance-based model (***Destexhe et al., 1994***), the average of the peak PSC can be shown to be proportional to the total number of released neurotransmitters (***Katz and Miledi, 1965***):

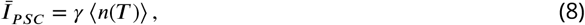

where *T* is the duration of the action potential (~ 1*ms*) and *γ* is a factor that only depends on the properties of the postsynaptic neuron. As our focus is on the role of neurotransmitter release dynamics in synaptic transmission, *γ* can be regarded as a constant, and ⟨*n*(*T*)⟩ and 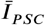 can be used interchangeably. Equations (3), (5) and (8) relate the presynaptic action potential to the postsynaptic current response and thus complete our framework for synaptic transmission.

The present theory was motivated by the experimental setups that are capable of probing synaptic transmission at the single-synapse level. The single-synapse level of description does not limit the theory to single-input synapses; rather, it opens up access to information that is otherwise lost to averaging in the bulk-level description. A postsynaptic neuron usually receives inputs from many synaptic connections, and the cellular response is an integration of these inputs. In the case of multiple synaptic inputs, the theory applies to each synapse separately and will yield an individual set of microscopic parameters for each synapse. The postsynaptic response can be calculated as a nonlinear summation of these inputs. The analytic expressions presented above can be readily applied to integrated multiple synaptic inputs when the molecular features of the presynaptic and postsynaptic sides are similar across the synapses, as in the case of the synapses that originate from the same axon and connect to nearby dendritic regions of a postsynaptic neuron (***Branco and Staras, 2009***). The generalization of the theory to elucidate the effects of dendritic integration of large heterogeneous synaptic inputs is a subject of future work. Detailed derivations of Eqs. (2)–(8) are given in Appendix 1.

To validate the developed analytic theory, we first compare its predictions to data generated through numerical simulations of the kinetic scheme in Eq. (1). A simple least squares fit reliably recovers input parameters of the simulations (Appendix 2 Figure 1). Next, we test the robustness of the theory by comparing it to modified simulations, in which deviations from the assumptions underlying Eqs. (2)–(5) are introduced. The modified simulations incorporate (i) the finite-capacity effect of RRP and (ii) heterogeneity of [*Ca*^2+^] among different release sites. For deviations within physiological range, the analytic expressions still reliably recover input parameters (Appendix 2 Figure 2). Details of the simulations are given in Appendix 2.

### Application of the theory to experimental data

Analytic expressions for measurable quantities enable direct application of the theory to experimental data.

A fit of the peak release rate vs. [*Ca*^2+^] with Eqs. (5) and (6) was performed for different synapses to extract a set of parameters {Δ*G*^‡^, *Q*^‡^, *k*_0_} for each synapse. These parameters were then used to rescale the peak release rate and calcium concentration to get the dimensionless quantities *r* and *c* that appear in Eq. (7). The experimental data used in the fit included *in vivo* measurements on (i) the Calyx of Held, a large synapse (diameter ~ 20*μm*) in the auditory central nervous system, at different developmental stages (***Schneggenburger and Neher, 2000; Lou et al., 2005; Bollmann et al., 2000***); (ii) parallel fiber - molecular layer interneuron (PF-MLI), a small synapse (~ 1*μm*) in the cerebellum (***Miki et al., 2018***); (iii) the photoreceptor synapse (***Duncan et al., 2010***); and (iv) two *in vitro* measurements (***Kyoung et al., 2011; Diao et al., 2012***). Figure 2A demonstrates that the data from these different synapses collapse on a single curve given by Eq. (7), consistent with the prediction of the theory. Remarkably, while release rates naturally have huge variation among different synapses, our theory reveals that they all can be brought into a compact, universal form. The universal collapse is an indication that synaptic transmission in different synapses is governed by common physical principles and that these principles are captured by the present theory. Variability across synapses on the molecular level is captured through the distinct sets {Δ*G*^‡^, *Q*^‡^, *k*_0_} for each synapse. Due to its mechanistic nature, the present theory can be further tested by independently measuring the parameters {Δ*G*^‡^, *Q*^‡^, *k*_0_} through single-molecule experiments and the postsynaptic response through electrophysiological recording experiments, which offers an advantage over the previous work based on phenomenological formulas (***Kochubey et al., 2011***). Notably, the generality of Eq. (7) spans beyond the context of synaptic transmission: the same scaling has appeared in another, seemingly unrelated, instance of biological membrane fusion – infection of a cell by an enveloped virus (***Zhang and Dudko, 2015***).

**Figure 2.**
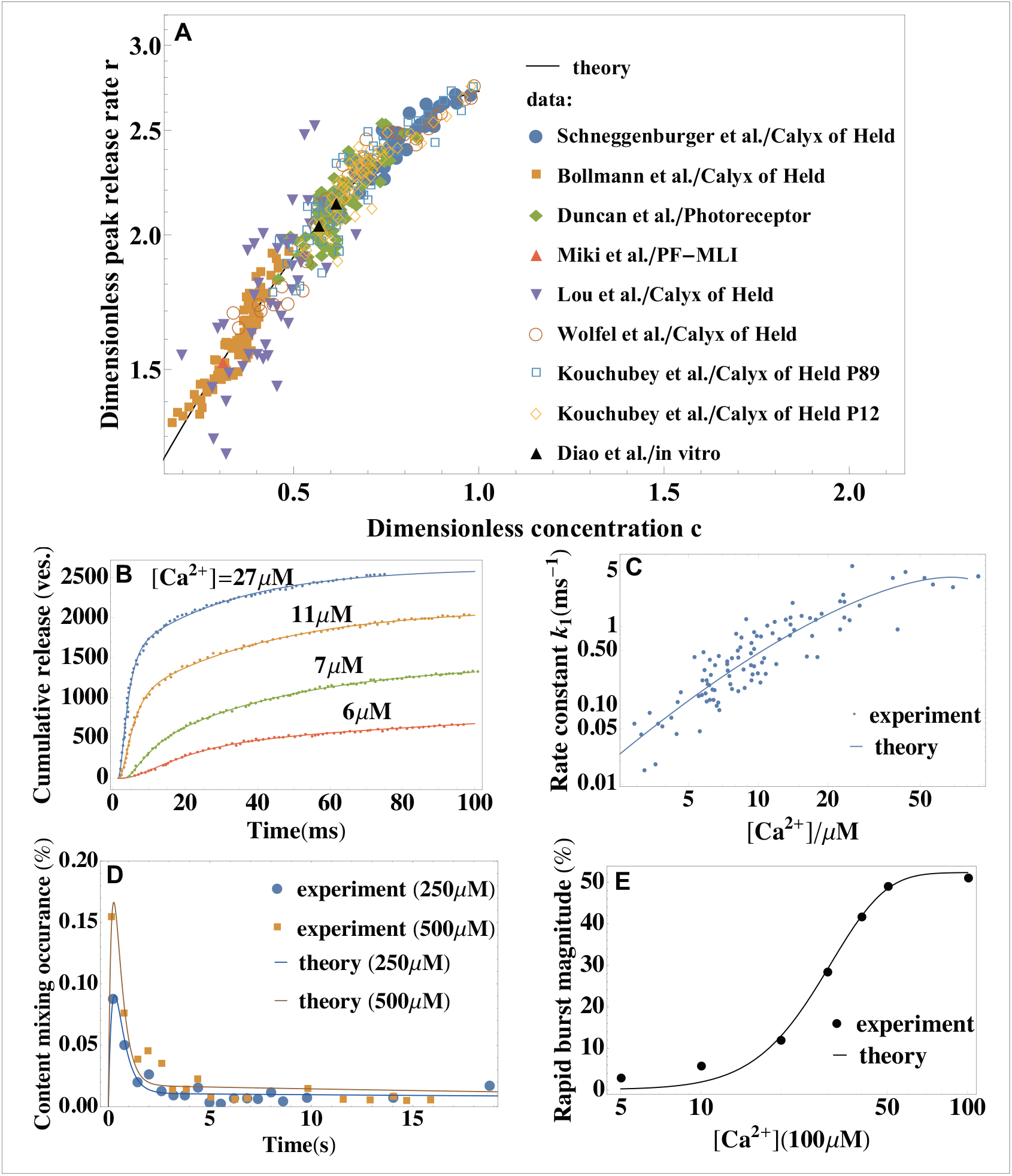
Application of the theory to experiments: extracting parameters of synaptic fusion machinery and verifying universality. (A) Dimensionless peak release rate as a function of dimensionless calcium concentration for a variety of synapses (***Schneggenburger and Neher, 2000; Lou et al., 2005; Bollmann et al., 2000; Miki et al., 2018; Duncan et al., 2010; Diao et al., 2012; Kochubey et al., 2009; Wölfel et al., 2007***). The data collapse onto a universal curve, Eq. (7), highlighting universality in synaptic transmission. Variability across synapses is captured through distinct sets of parameters extracted from the fit with Eq. (6) (Appendix 3 Table 2). (B) Cumulative release from *in vivo* experiments (***Wölfel et al., 2007***) (symbols) and a fit with Eq. (3) (lines) for different calcium concentrations. (C) *Ca*^2+^-dependent rate constant of SNARE conformational transition from *in vivo* experiments (***Wölfel et al., 2007; Kochubey et al., 2009***) and a fit with Eq. (5). (D) Content mixing occurrence from *in vitro* experiments (***Diao et al., 2012***) and a fit with Eq. (2). (E) Rapid burst magnitude from *in vitro* experiments (***Kyoung et al., 2011***) and Eq. (3). Parameters are shown in Appendix 3.

The utility of the theory as a tool for extracting microscopic parameters of synaptic fusion machinery is further illustrated in Fig. 2B-E. A fit of *in vivo* data for cumulative release at different [*Ca*^2+^] (***Wölfel et al., 2007***) with Eq. (3) extracts the rate of SNARE conformational transition, *k*_1_([*Ca*^2+^]) (Fig. 2B). A fit of the rate with Eq. (5) extracts the SNARE activation barrier and rate at reference concentration [*Ca*^2+^]_0_ (Fig. 2B). Fits of *in vitro* data (***Kyoung et al., 2011; Diao et al., 2012***) with Eqs. (2) and (3) are shown in Fig. 2D and E. In Fig. 2D, the content mixing occurrence, defined in (***Kyoung et al., 2011***) as the average release rate normalized by the total number of vesicles, 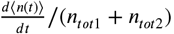, is fitted with Eq. (2). In Fig. 2E, the rapid burst magnitude, defined in (***Diao et al., 2012***) as the ratio of the numbers of vesicles fused within the first 1*s* and within 50*s* after calcium trigger, ⟨*n*(*t* = 1*s*)⟩ /⟨*n*(*t* = 50*s*)⟩, is fitted with Eq. (3).

Details of the fitting procedure and the parameter values extracted from the fits in Fig. 2 are given in Appendix 3.

### Linking molecular mechanisms to synaptic function

#### Short-term plasticity

Synaptic plasticity, or the ability of synapses to strengthen or weaken over time depending on the history of their activity, underlies learning and memory (***Regehr, 2012; Bailey et al., 2015***). A measure of synaptic strength is the peak of the post-synaptic current, which is proportional to cumulative release (Eq. (8)). The short-term change (< 1min) in synaptic strength, known as short-term plasticity (***Regehr, 2012***), can be estimated through the ratio of the cumulative release for two consecutive action potentials of width *T* (typically *T* ~ 1/*k*_1_ ≪ 1/*k*_2_) separated by interpulse interval *τ*_*int*_. Whether the synapse weakens or strengthens is determined by the relative timescales of two competing factors: residual *Ca*^2+^ removal, *τ*_*Ca*_, and RRP vesicle replenishment, *τ*_*RRP*_. On the one hand, residual calcium at the presynaptic terminal results in a higher calcium concentration during the subsequent action potential, [*Ca*^2+^]_*f*_ > [*Ca*^2+^]_*i*_, which tends to strengthen the synapse. On the other hand, RRP vesicle depletion results in *n*_*tot*1,*i*_ > *n*_*tot*1,*f*_, which tends to weaken the synapse. Assuming first-order kinetics of the calcium concentration and RRP vesicle replenishment, the change in synaptic strength can be obtained from Eq. (3) as (Appendix 1)

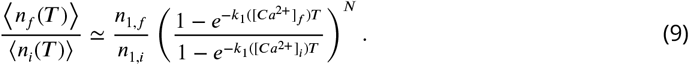

Equation (9) predicts that, for fixed interpulse interval *τ*_*int*_, the synapse will exhibit short-term facilitation or short-term depression depending on the ratio of the timescales, *τ*_*Ca*_/*τ*_*RRP*_, as illustrated in Fig. 3A (***Tank et al., 1995***). Furthermore, Eq. (9) shows that a given synapse may exhibit multiple forms of short-term plasticity when the interpulse interval *τ*_*int*_ is varied (Fig. 3A). Such coexistence of multiple forms of plasticity has been observed experimentally (***Regehr, 2012***).

**Figure 3.**
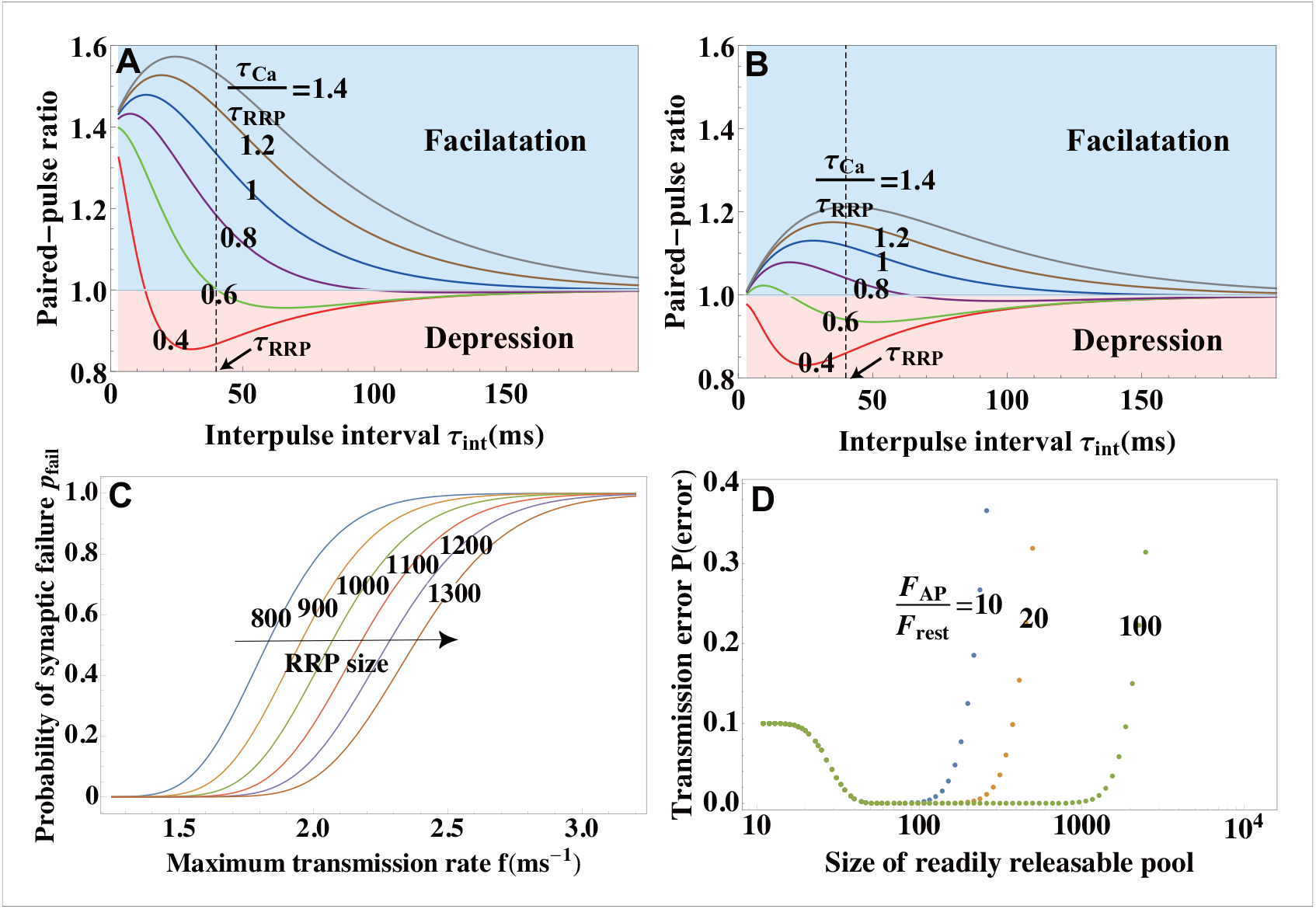
Functional implications of the theory. (A,B) Paired-pulse ratio of peak postsynaptic currents as a measure of short-term plasticity (the change in synaptic strength), assessed through the ratio of the cumulative release for two consecutive action potentials separated by interpulse interval *τ*_*int*_, Eq. (9). Synapses exhibit short-term facilitation or depression depending on the relative timescales of the replenishment of the readily releasable pool, *τ*_*RRP*_, and removal of residual calcium, *τ*_*Ca*_. A given synapse can exhibit multiple forms of short-term plasticity as the time interval *τ*_*int*_ is varied. Higher *Ca*^2+^-sensitivity of SNAREs (defined in Appendix 3; in A, it is 3 times that in B) results in a larger dynamic range of short-term plasticity. (C) Trade-off between the maximum transmission rate *f* = 1/*T* and fidelity 1 − *p*_*fail*_ from Eq. (10) for different RRP sizes. (D) Synaptic efficacy, 1 − *P*(*error*), from Eq. (12). The plateau around the optimal synaptic strength (Eq. (13)) indicates that no fine-tuning is required for near-optimal transmission of large synapses. Higher *Ca*^2+^-sensitivity *F*_*AP*_ / *F*_*rest*_ results in broader plateau for near-optimal performance.

A notable feature of Eq. (9) is the existence of an optimal value of interpulse interval at which facilitation (at large *τ*_*Ca*_/*τ*_*RRP*_) or depression (at small *τ*_*Ca*_/*τ*_*RRP*_) of synaptic transmission is maximal (Fig. 3A). Such optimality becomes less pronounced at intermediate values of *τ*_*Ca*_/*τ*_*RRP*_ where the synapse exhibits both facilitation and depression (see the curve at *τ*_*Ca*_/*τ*_*RRP*_ = 0.6 in Fig. 3A), suggesting a more subtle role of short-term plasticity in transmitting transient signals (***Tsodyks and Markram, 1997; Fuhrmann et al., 2002***). We further find that a lower *Ca*^2+^-sensitivity of a synapse (in Fig. 3B it is 1/3 of that in Fig. 3A) results in a smaller response range, indicating that high *Ca*^2+^-sensitivity of synaptic fusion machinery is essential for the large dynamic range of short-term plasticity in synaptic transmission.

Finally, Eq. (9) reveals how differences among synapses on the molecular level manifest themselves as different short-term facilitation/depression modes (Appendix 1 Figure 2). The molecular-scale differences are captured through different sets of parameters {Δ*G*^‡^, *Q*^‡^, *k*_0_} and *τ*_*Ca*_ and can be due to different variants of synaptotagmin in SNAREs (***Hui et al., 2005***), different coupling mechanisms of SNAREs and the scaffolding proteins at release sites (***Gramlich and Klyachko, 2019***), or different types of *Ca*^2+^ buffering proteins present at the presynaptic terminal (***Schwaller, 2020***). This finding suggests that the diversity of the molecular machinery for vesicle fusion enables the diversity of synaptic function in short-term plasticity.

#### Transmission rate vs. fidelity

An important characteristic of neuronal communication is fidelity of synaptic transmission. Two measures of fidelity can be considered at the single-synapse level for different types of synapses. The probability of spike transmission is a natural measure of fidelity for giant synapses in sensory systems ***Borst and Soria van Hoeve*** (***2012***) and neuromuscular junctions. The probability of a postsynaptic voltage/current response, beyond the noise level, to a presynaptic spike is a measure of fidelity for small synapses in the central nervous system (CNS) (***Dobrunz and Stevens, 1997***). The probabilistic nature of release mechanisms at synapses is a common origin of synaptic failure (***Allen and Stevens, 1994***).

Although the two definitions of fidelity apply to different types of synapses, the present theory allows for a unifying treatment of both phenomena. We assume that the desired postsynaptic response – a postsynaptic spike or a postsynaptic current beyond the noise level – is generated only if the number of released vesicles in response to an action potential exceeds some threshold *M*. The value of *M* depends on the density of ion channels on the postsynaptic membrane and the excitability of the postsynaptic neuron (***Biederer et al., 2017***). For both types of the postsynaptic response, the probability that the synaptic transmission fails is then obtained from the probability *P*{*n*(*t*) = *m*} that *m* vesicles fuse by time *t* as

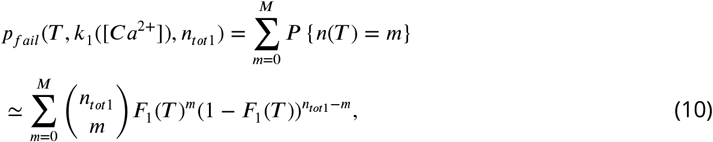

where 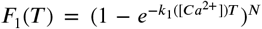. Since the presynaptic neuron cannot generate a second spike during time [0, *T*], *f* ≡ 1/*T* represents the maximum transmission rate. Equation (10) predicts that a higher transmission rate *f* results in a higher probability of transmission failure *p*_*fail*_ and thus lower fidelity (1 − *p*_*fail*_). This trade-off between the maximum rate and fidelity in synaptic transmission is shown in Fig. 3C. Consistent with intuitive expectation, Eq. (10) further predicts that, for a given maximum transmission rate, the probability of transmission failure can be reduced by increasing the RRP size *n*_*tot*1_ and/or SNARE conformational rate *k*_1_([*Ca*^2+^]) (Fig. 3C).

Equation (10) allows us to make a quantitative statement regarding the fidelity of synapses of different sizes. Faithful spike transmission implies that the threshold *M* for postsynaptic response is smaller than the average cumulative release, *M* < ⟨*n*(*T*)⟩ = *n*_*tot*1_*F*_1_(*T*). Then, by the Chernoff bound for Eq. (10) (***Vershynin, 2018***),

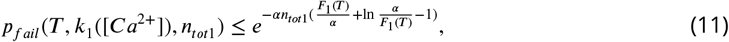

where *a* ≡ *M*/*n*_*tot*1_. Because both *M* and *n*_*tot*1_ scale linearly with the area of synaptic junctions (***Miki et al., 2017; Nakamura et al., 2015***), it is reasonable to assume that *a* = *M*/*n*_*tot*1_ < *F*_1_(*T*) is kept at an approximately constant level. Since *F*_1_(*T*)/*α* + ln(*α*/*F*_1_(*T*)) − 1 > 0, the probability of synaptic failure decreases exponentially as the RRP size *n*_*tot*1_ increases. Thus, it follows from Eq. (11) that larger synapses tend to be significantly more reliable, i.e., have an exponentially smaller probability to fail, than smaller synapses in transmitting signals (***Dobrunz and Stevens, 1997***).

#### Synaptic efficacy

Equations (10)–(11) show that synaptic strength can be increased, i.e. failure suppressed, by increasing the RRP size or decreasing the threshold for eliciting postsynaptic response. However, a high synaptic strength increases the probability of an error read, i.e. a postsynaptic response generated without a presynaptic spike. We will now establish the condition for the optimal synaptic strength through the balance of probabilities of failure (no postsynaptic response to an action potential) and error read (postsynaptic response in the absence of an action potential). Let [*Ca*^2+^]_*r st*_ and [*Ca*^2+^]_*AP*_ be the calcium concentrations at rest and during the action potential and *q* the probability of firing an action potential by the presynaptic neuron. The total probability of transmission error is

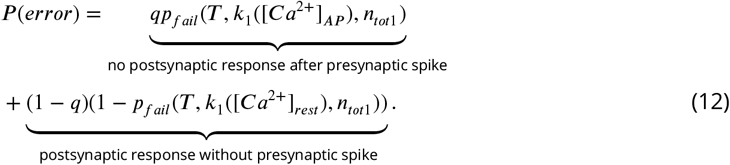

Here, we assume that the long-term (minutes to days) change in synaptic strength, known as long-term plasticity, occurs through the presynaptic mechanisms and is predominantly due changes in the RRP size, *n*_*tot*1_, which has been shown to be regulated through retrograde signaling according to the threshold *M* on the postsynaptic side (***Yang and Calakos, 2013; Mayford et al., 2012; Bailey et al., 2015***). Synaptic efficacy, 1 − *P*(*error*), measures the ability of the synapse to faithfully transmit signal. The optimal RRP size is obtained by minimizing the transmission error in Eq. (12):

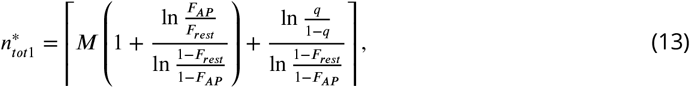

where ⌈*x*⌉ denotes ceiling, i.e. the smallest integer greater than or equal to *x*, and 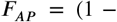 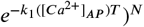 and 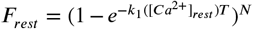. are the fusion probabilities during the action potential and at rest. Equation (13) predicts that, as the synapse is stimulated more frequently (*q* increases), a larger RRP size is needed for the optimal performance, i.e. the optimal RRP size and hence the optimal synaptic strength increase, resulting in long-term potentiation on the presynaptic side.

How far can the RRP size deviate from its optimal value without a significant loss of synaptic efficacy? The range of RRP sizes for near-optimal performance can be estimated through the Chernoff bound for Eq. (12):

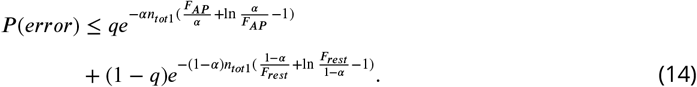

According to Eq. (14), for synapses that are large (*n*_*tot*1_ ≫ 1) and sufficiently sensitive to *Ca*^2+^ (*F*_*AP*_ /*F*_*rest*_ ≫ 1), the error probability is exponentially small and thus insensitive to changes in the RRP size *n*_*tot*1_. Specifically, the near-optimal range for *n*_*tot*1_ can be estimated from *F*_*rest*_ ≲ *α* ≲ *F*_*AP*_ to be *M*/*F*_AP_ ≲ *n*_*tot*1_ ≲ *M*/*F*_*rest*_. Since 1/*F*_*AP*_ ≪ 1/*F*_*rest*_, this range is broad, indicating that the synapses do not need to fine-tune their RRP size in order to maintain near-optimal transmission. This robustness in synaptic transmission is illustrated in Fig. 3D.

## Discussion

The capacity of neurons to transmit information through synapses rapidly and precisely is the key to our ability to feel, think, or perform actions. Despite the challenge posed for experimental studies by the ultrashort timescale of synaptic transmission, a number of recent experiments *in vivo* (***Schneggenburger and Neher, 2000; Lou et al., 2005; Bollmann et al., 2000; Miki et al., 2018; Duncan et al., 2010; Kochubey et al., 2009; Wölfel et al., 2007***) and in reconstituted systems (***Kyoung et al., 2011; Diao et al., 2012***) demonstrated the ability to probe the kinetics of synaptic transmission at the single-synapse level. By design, these experiments generate pre-averaged data that encode unprecedented information on the molecular mechanisms of synaptic function, which is lost in the data that are averaged over multiple heterogeneous synaptic inputs. However, decoding this information requires a quantitative framework that would link the quantities that are measured in the experiments to the microscopic parameters of the synaptic release machinery. Here, we presented a statistical-mechanical theory that establishes these links.

### Analytic theory for synaptic transmission

Our theory casts the synaptic fusion scenarios observed in different experimental setups into a unifying kinetic scheme. Each step in this scheme has its mechanistic origin in the context of a given experimental setup. In the context of *in vivo* experiments, distinct vesicle pool dynamics are taken into account (***Alabi and Tsien, 2012; Yang and Calakos, 2013; Kaeser and Regehr, 2017***) to quantitatively explain the different timescales observed in the vesicle release dynamics (***Kaeser and Regehr, 2014; Neher, 2017; Rozov et al., 2019***): vesicles from the readily releasable pool (RRP) fuse readily once the critical number of SNARE complexes undergo conformational transitions upon *Ca*^2+^ influx (fast step), while the reserve pool supplies vesicles to the RRP (slow step). In the context of *in vitro* experiments, different timescales in vesicle release dynamics are due to the observed distinct states of docked vesicles (***Diao et al., 2012; Kweon et al., 2017; Gipson et al., 2017***): the vesicles that are in a point contact with the membrane fuse readily upon *Ca*^2+^-triggered SNARE conformational transition (fast step), thus mimicking RRP vesicles *in vivo*, while the vesicles that are in an extended contact become trapped in a hemifusion diaphragm state prior to fusing with the membrane (slow step). Although the role of these distinct docked states *in vivo* is still under debate (***Neher and Brose, 2018; Brunger et al., 2018***), the realization that both of the fusion scenarios can in fact be mapped onto the same kinetic scheme allowed us to capture these scenarios through a unifying theory. The fact that each fusion step in the kinetic scheme has a concrete mechanistic interpretation makes the theory directly predictive in both *in vitro* and *in vivo* experiments.

The developed analytic theory yields closed-form expressions for measurable quantities. The expressions enable the extraction of microscopic parameters of synaptic release machinery through a simple fit to experimental data. The calculated measurable quantities include: (i) cumulative release (***Miki, 2019***), which quantifies the number of vesicles fused during a given time interval following the action potential, (ii) temporal profile of the release rate, which measures the rate of change in the number of fused vesicles, (iii) peak release rate, which is a measure of sensitivity of a synapse to the trigger, and (iv) the calcium-dependent rate of SNARE conformational change. A least-squares fit of data with these expressions yields the activation energy barrier and rate constant for SNARE conformational change at any calcium concentration of interest, the critical number of SNAREs necessary for fusion, and the sizes of the readily releasable and reserve vesicle pools.

The theory is developed at the single-synapse level and is applicable both to giant synapses with many active zones in sensory systems (***Borst and Soria van Hoeve, 2012***) and to small synapses with few active zones in the brain (***Harris and Weinberg, 2012***). For a postsynaptic neuron that receives inputs from heterogeneous synaptic connections, the theory applies to each individual synapse and will provide a quantitative framework for studying how heterogeneity among the synapses influences their integrated effect on the postsynaptic neuron. We validate the theory through two sets of numerical simulations. First, we demonstrate that the derived analytic expressions reliably recover the (known) parameters of the simulations of the underlying kinetic scheme. Next, we perform simulations with deliberately introduced, biologically relevant deviations from the assumptions of the theory, namely the finite-capacity effect of the RRP (***Biederer et al., 2017***) and heterogeneity of calcium concentration among different release sites ***Gramlich and Klyachko*** (***2019***). For deviations within physiological range, the expressions still reliably recover input parameters, demonstrating the robustness of the analytic theory.

### Conformational transition of SNARE proteins

The rate-limiting step in the initiation of fusion of the synaptic vesicles that are docked on the presynaptic membrane is the conformational transition of the critical number of SNARE proteins tethering the vesicles to the membrane (***Kaeser and Regehr, 2014***). We derived the calcium-dependence of the SNARE conformational rate from the classical reaction-rate theory (***Kramers, 1940***) which was generalized to include an external trigger – calcium concentration. The resulting analytic expression reveals that the SNARE conformational rate, and hence the vesicle release rate and the peak of the release rate, are all exponentially sensitive to calcium concentration. This result provides a quantitative explanation for the observed remarkable synchrony of synaptic vesicle fusion: since the rising of calcium concentration after an action potential occurs on a microsecond timescale and is thus essentially instantaneous on the timescale of synaptic release, the exponential sensitivity of the release rate to this nearly-instantaneous trigger ensures an ultra-rapid initiation of vesicle fusion upon calcium influx; likewise, the exponential sensitivity of the release rate to calcium ensures that the fusion process terminates rapidly upon calcium depletion (***Brunger et al., 2018***).

### Critical number of SNAREs for vesicle fusion

The theory further reveals how the kinetics of vesicle fusion are affected by the critical number of SNAREs per vesicle. Given the lack of general consensus (***Südhof, 2013; Sinha et al., 2011; van den Bogaart and Jahn, 2011; Brunger et al., 2018***), the theory makes no assumptions about the specific number of SNAREs necessary for fusion, and the number itself can serve as a free parameter when sufficient data is available for a robust fit. Interestingly, however, the theory suggests that *N* = 2 SNAREs per vesicle provide the optimal balance between stability and precision of release dynamics. Indeed, on the one hand, in the presence of a single SNARE, the high values and an exponentially-steep temporal dependence of the release rate makes the rate very sensitive to sub-millisecond calcium fluctuations, and thus a very fine tuning of the calcium concentration would be necessary to prevent instability of the fusion process. On the other hand, the values of *N* greater than 2 lead to longer delays in the peak of the release rate following an action potential, thus reducing the temporal precision of vesicle release. Furthermore, a least-squares fit of the release rate from the experiment (***Kochubey et al., 2009***) with the theory at different values of *N* reveals that *N* = 2 indeed results in the smallest fitting errors for all calcium concentrations. The generality of this result can be determined as more data on the release dynamics for different synapses becomes available.

### Universality vs. specificity in synaptic transmission

The fact that, in all chemical synapses, the delay time from the action potential triggering to vesicle fusion is determined by the conformational transition of preassembled SNARE complexes, and that the conformational transition itself occurs through a single rate-limited step, suggests possible universality in synaptic transmission across different synapses despite their structural and kinetic diversity. Our theory made this intuition precise through a nondimensionalized scaling relationship between the peak release rate and calcium concentration (Eq. (7)), which is predicted to hold for all synapses irrespective of their variability on the molecular level. In statistical physics, the significance of universality is that it indicates that the observed phenomenon (here, synaptic transmission) realized in different systems is governed by common physical principles that transcend the details of particular systems.

Once validated through a comparison with numerical simulations, the universal relation was tested using published experimental data on a variety of synapses, including *in vivo* measurements on the Calyx of Held (an approximately 20*μm*-diameter synapse in the auditory central nervous system) studied at different developmental stages, parallel fiber-molecular layer interneuron (a 1*μm*-diameter synapse in the cerebellum) and the photoreceptor synapse, as well as a reconstituted system. Despite more than an order of magnitude difference in the size and nearly three orders of magnitude variation in the dynamic range of these synapses, the data for the sensitivity of the synapses to the trigger collapsed onto a universal curve, as predicted by the theory. The collapse serves as an evidence that the established scaling of the normalized peak release *r* with calcium concentration *c*, *r* = exp [1 − (1 − *c*)^3/2^], is indeed universal across different synapses. At the same time, the unique properties of each synapse are captured by the theory through the distinct sets of parameters of their molecular machinery: the critical number of SNAREs, their kinetic and energetic characteristics, and the sizes of the vesicle pools. The practical value of the theory as a tool for extracting microscopic parameters of synapses was further illustrated by fitting *in vivo* and *in vitro* data for cumulative release and for the average release rate at different calcium concentrations. Compared to previous work based on phenomenological formulas (***Kochubey et al., 2011***), the mechanistic nature of the present theory makes its predictions testable through single-molecule manipulation experiments (***Gao et al., 2012; Oelkers et al., 2016***), which can provide independent estimates for the microscopic parameters of synaptic fusion machinery.

### From molecules to synaptic function

We applied the theory to establish connections between the molecular constituents of synapses and synaptic function. The theory provides an analytic expression for the short-term change in synaptic strength, or short-term plasticity. The expression quantitatively describes how short-term facilitation, short-term depression, or coexistence of multiple forms of plasticity in a given synapse emerge from the interplay between two molecular-scale processes – residual calcium removal and replenishment of the readily releasable pool. In contrast to phenomenological models of short-term plasticity (***Tsodyks and Markram, 1997; Fuhrmann et al., 2002; Rosenbaum et al., 2012***), the mechanistic nature of the present theory allows us to explore how temporal filtering of synaptic transmission is affected by calcium-sensitivity of synaptic fusion machinery, as well as how diverse short-term facilitation/depression modes emerge from the diversity of the molecular constituents. Furthermore, while one intuitively expects that there must be a tradeoff between the maximum transmission rate and fidelity of a synapse, our theory turns this intuition into a quantitative relation, which quantifies how the probability of transmission failure can be reduced through changes of the size of the readily releasable pool and SNARE conformational rate. The tradeoff relation further shows that the probability of synaptic failure decreases exponentially with increasing the synapse size, which makes large synapses significantly more reliable than small synapses in transmitting signals. We established a condition for the maximal synaptic efficacy, at which the combined probability of failure (lack of postsynaptic response to an action potential) and error read (a response in the absence of an action potential) is minimized. Interestingly, we find that, for large synapses, the parameter range of near-optimal performance is broad, indicating that no fine tuning is needed for these synapses to maintain near-optimal transmission. This finding may also be relevant to small synapses: although a small size of their individual RRPs makes them less reliable in transmitting signals individually, trans-synaptic interactions that couple many nearby small synapses may result in a large “effective” RRP (***Bailey et al., 2015***) and thus enable small synapses to collectively maintain near-optimal transmission without fine-tuning. Altogether, the results of the theory provide a quantitative basis for the notion that the molecular-level properties of synapses are not merely details but are crucial determinants of the computational and information-processing synaptic functions (***Südhof, 2013***).

Other biological processes, including infection by enveloped viruses, fertilization, skeletal muscle formation, carcinogenesis, intracellular trafficking, and secretion, have features that are very similar to those in synaptic transmission, despite the bewildering number and structural diversity of the molecular constituents involved (***Harrison (2017***)). These processes occur through membrane fusion that (i) requires overcoming high energy barriers, (ii) is controlled by proteins that undergo a conformational transition once exposed to a trigger, (iii) is facilitated by the energy released during this transition, which reduces the fusion timescale by orders of magnitude. The theory presented here can be generalized to encompass these processes while engaging with the diversity of specific systems. The mapping from molecular mechanisms to cellular function, provided by the present theory, is a step toward a more complete framework that would bridge mechanisms with function at the multicellular scale (e.g. neuronal circuits and tissues) and further at the scale of an organism.

## Materials and Methods

Details of the derivations for analytical results, simulation methods and fitting procedures are described in Appendices 1, 2 and 3.

## Acknowledgments

We are grateful to Bill Bialek for insightful discussions, and Tynan Kennedy and Palka Puri for critical comments on the manuscript. This research was supported by the National Science Foundation Grant No. MCB-1411884.

## Appendix 1 Derivations for analytic results

### Synaptic fusion dynamics and fusion times

As discussed in the main text, even though the reaction schemes for calcium-triggered synaptic release *in vivo* and *in vitro* differ in the interpretation of the individual states, their mathematical equivalence enables the treatment through a unified theory. Independent of the details of the experimental situation, synaptic fusion generally involves a “fast” pool and a “slow” pool of synaptic vesicles (Fig. 1 in the main text).

We assume that fusion of a vesicle from the fast pool requires a conformational change of each of the *N* SNARE complexes attached to it and that the SNAREs undergo their conformational changes independently with the corresponding times *τ*_1_, *τ*_2_…*τ*_*N*_. Since the conformational change of a SNARE complex is dominated by a single barrier (***Hui et al., 2005***), *τ*_*i*_ satisfies the exponential distribution

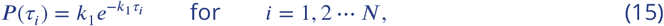

where *k*_1_ ≡ *k*_1_([*Ca*^2+^]) is the calcium-dependent rate constant for the conformational change of each SNARE complex. The fusion time for a vesicle in the fast pool, *T*_1_, is therefore determined by the largest value of *τ*_*i*_. For an action potential, and the rise of calcium concentration, triggered at time *t* = 0, the probability that fusion has occurred by time *t* is

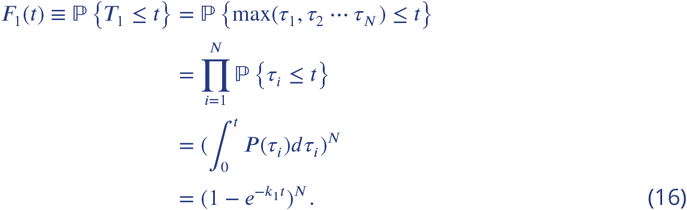

Eq. (16) is the cumulative distribution for the fusion times *T*_1_ of vesicles in the fast pool. The corresponding probability density function is

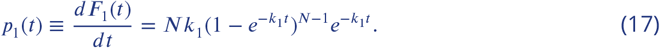

The fusion process for vesicles in the slow pool consists of two sequential steps: conformation changes of *N* SNAREs with rate constant *k*_1_ and a slow reaction step with rate constant *k*_2_. *In vivo*, the slow step corresponds to the replenishment of the readily releasable pool (RRP) with vesicles from the reserve pool through their docking and priming. *In vitro*, the slow step corresponds to the escape from the metastable “hemifusion diaphragm” state of the vesicle. Thus, the fusion time for vesicles in the slow pool is 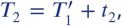, where 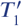 has the same probability distribution as *T*_1_ (Eqs. (16) and (17)) and *t*_2_ satisfies the exponential distribution with parameter *k*_2_. Since 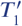 and *t*_2_ are mutually independent, the probability density distribution for *T*_2_ is the following convolution:

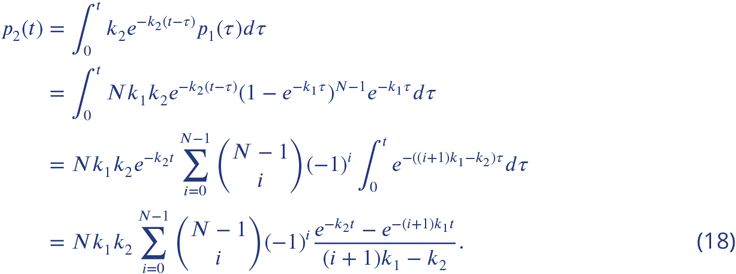

We use 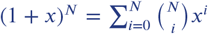 in the third line. The cumulative distribution for *T*_2_ is obtained by integration of Eq. (18):

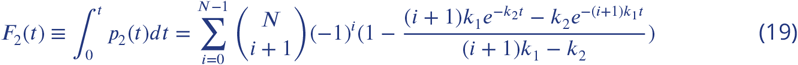

### Synaptic release statistics and the average release rate

The cumulative distributions for fusion time, Eqs. (16) and (19), allow us to calculate the probability distribution for the number of vesicles from the fast and slow pathways, *n*_1_(*t*) and *n*_2_(*t*), that have fused by time *t*. Assuming that fusion of vesicles within each pool is independent and random, we have

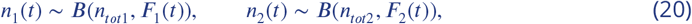

where *B*(*n, p*) is binomial distribution with parameters *n* and *p*, and *n*_*tot*1_ and *n*_*tot*2_ are the sizes of the fast and slow pools. Furthermore, assuming that vesicles in the fast and slow pools are released independently, the distribution of the total number of released vesicles, *n*(*t*) = *n*_1_(*t*) + *n*_2_(*t*), is found as the convolution of two binomial distributions:

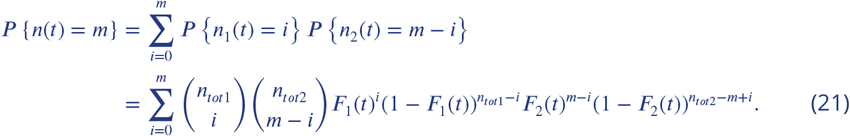

Equation (21) gives the probability that *m* vesicles fuse by time *t*. When the slow step *k*_2_ is neglected, *F*_2_(*t*) ~ 0, Eq. (21) reduces to the binomial distribution (***Malagon et al., 2016***): 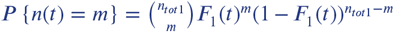. Using Eq. (21), the critical number of SNAREs, *N*, can be accurately determined by Bayesian model comparison from the release statistics.

The average cumulative release is then

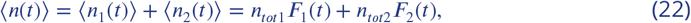

where *F*_1_(*t*) and *F*_2_(*t*) are given by Eq. (16) and (19). This is Eq. (3) in the main text.

The average release rate is obtained by differentiating Eq. (22):

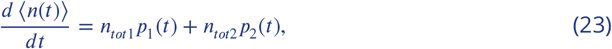

where *p*_1_(*t*) and *p*_2_(*t*) are given by Eqs. (17) and (18). This is Eq. (2) in the main text.

### Average release in various asymptotic regimes

Examining the asymptotic behavior of the exact solutions derived above yields approximate yet simple and accurate expressions for experimentally measurable quantities.

On the short timescale, *t* ≪ 1/*k*_1_ ≪ 1/*k*_2_, keeping the leading-order term *k*_1_*t* and neglecting all the terms related to *k*_2_*t* ≪ *k*_1_*t* in Eq. (23), we have

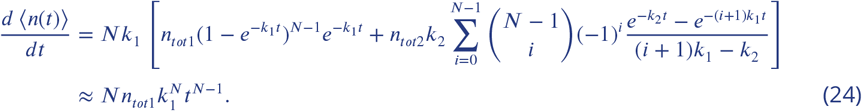

Equation (24) provides a means to test the kinetic scheme in Eq. (1) in the main text and distinguish it from the previously proposed scheme (***Bollmann et al., 2000; Miki et al., 2018; Schneggenburger and Neher, 2000***) that assumed that the binding of each calcium ion to a SNARE complex constitutes a separate rate-limiting step along the fusion reaction. Specifically, the schemes may be distinguished through the scaling behavior of 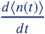 at short-times, *t* ≪ 1/*k*_1_. It can be shown that, for a general stochastic trajectory with sequential transitions that are characterized by rate constants 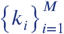, 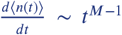 at times *t* ≪ 1/ max(*k*_*i*_). The scheme in Eq. (1) in the main text corresponds to *M* = *N* ~ 2. In contrast, the scheme based on individual calcium ion binding (***Bollmann et al., 2000; Miki et al., 2018; Schneggenburger and Neher, 2000***) corresponds to *M* ≥ 4. The time resolution of the current experiments *t*_*c*_ ~ 1/*k*_1_ ~ 1*ms* may not be sufficient to resolve these different behaviors, but future experiments may make this test possible.

On the intermediate timescale 1/*k*_1_ ≪ *t* ≪ 1/*k*_2_, we have *F*_1_(*t*) ≈ 1 and thus

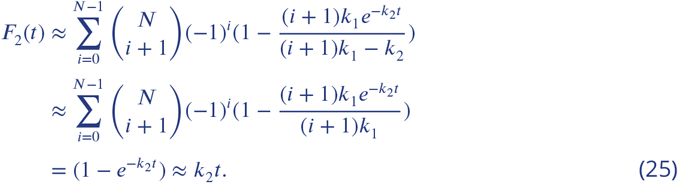

Therefore, Eq. (22) becomes

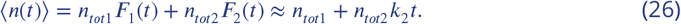

On the long timescale *t* ≫ 1/*k*_2_ ≫ 1/*k*_1_, we have *F*_1_(*t*) ≈ 1 and 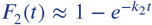 (see Eq. (25)), and thus

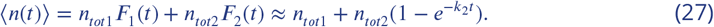

### Peak release rate

The time *t*_*max*_ at which the release rate is maximal can be obtained by setting the derivative of the average release rate in Eq. (23) to be zero:

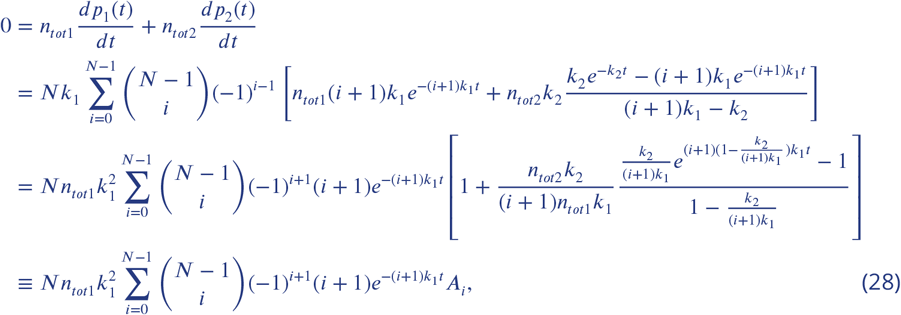

where we used 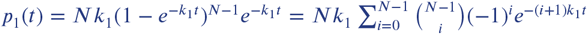 in the second line and introduced *A*_*i*_ in the last line for convenience. The time *t*_*max*_ is the solution of Eq. (28), which is a transcendental equation that is challenging to solve for *N* ≥ 2. In practice, *k*_2_/*k*_1_ ≪ 1, which allows us to expand *A*_*i*_ up to the first order of *k*_2_/*k*_1_:

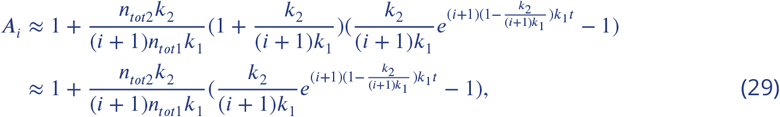

where high order terms are dropped in the second line. After dropping the constant factor 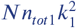, Eq. (28) becomes

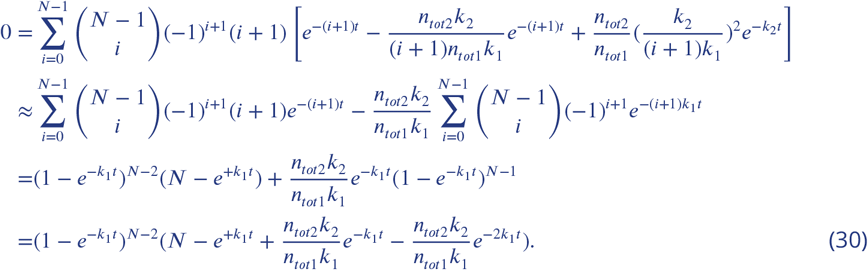

In the second line, the last term is dropped due to 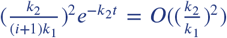. In the third line, we used 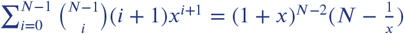 for *N* ≥ 2 and 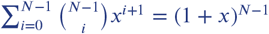.

Now let 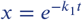 and 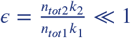. The equation for *t*_*max*_ up to order *O*(*ϵ*) becomes

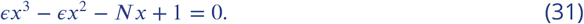

Assuming *x* = *x*_0_(1 + *ϵx*_1_ + *O*(*ϵ*^2^)) and comparing the coefficients of *ϵ*^0^ and *ϵ*^1^ in Eq. (31), we find: 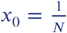 and 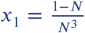. Therefore,

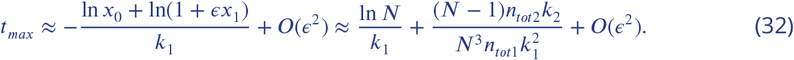

With 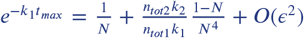, the peak release rate, up to the first order of 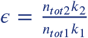, is

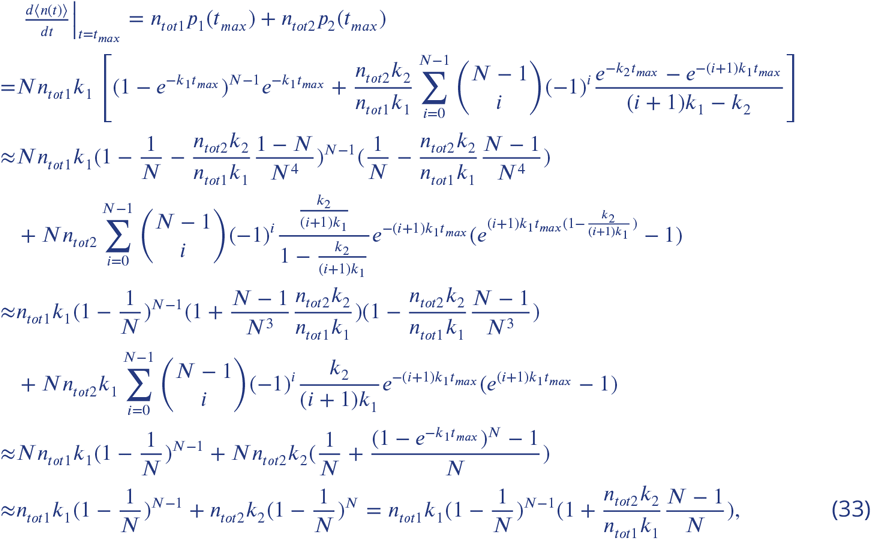

which is Eq. (4) in the main text.

### Calcium-dependent rate constant ***k***_*1*_([*Ca*^2+^])

Analogous to the expression for the *pH*-dependent Gibbs free energy of a protein (***Schaefer et al., 1997***), the calcium-dependent free energy can be written as

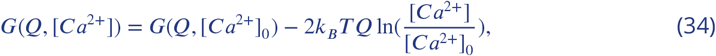

where *Q* is the average occupancy (relative to the reference value) of calcium ions on a SNARE complex, and the factor of 2 reflects the charge of a calcium ion being twice the charge of a hydrogen ion. The generic free energy profile *G*(*Q,* [*Ca*^2+^]_0_) at reference calcium concentration [*Ca*^2+^]_0_ with a well and a barrier can be captured by a cubic polynomial (***Dudko et al., 2006***):

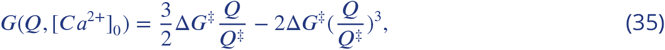

where Δ*G*^‡^ is the barrier height and *Q*^‡^ is the average occupancy number of calcium ions at the top of the barrier.

The reaction rate *k*_1_([*Ca*^2+^]) can then be derived from the Kramers formalism (***Kramers, 1940***) generalized to the presence of a bias field (***Dudko et al., 2006***). For given calcium concentration [*Ca*^2+^], the maximum of the free energy can be found from Eqs. (34) and (35) by solving *dG*(*Q,* [*Ca*^2+^])/*dQ* = 0:

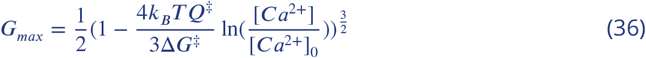

at

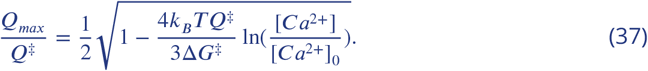

Due to symmetry in Eq. (35), the Kramers rate can be written as (***Kramers, 1940***)

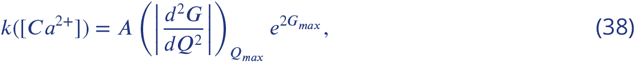

where 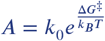 is a constant, independent of [*Ca*^2+^] and 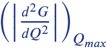 is the calcium-dependent curvature of *G*(*Q,* [*Ca*^2+^]) at *Q* = *Q*_*max*_.

Substituting Eqs. (36) and (34) into Eq. (38), we obtain the calcium-dependent reaction rate:

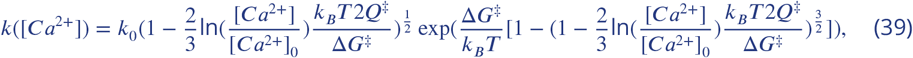

which is Eq. (5) in the main text. Appendix 1 Figure 1A shows the temporal profiles of the average release rate from the theory (Eqs. (2) and (5) in the main text) for different values of calcium concentration. Appendix 1 Figure 1B shows the average release rate for different values of the critical number of SNAREs, *N*.

**Appendix 1 Figure 1.**
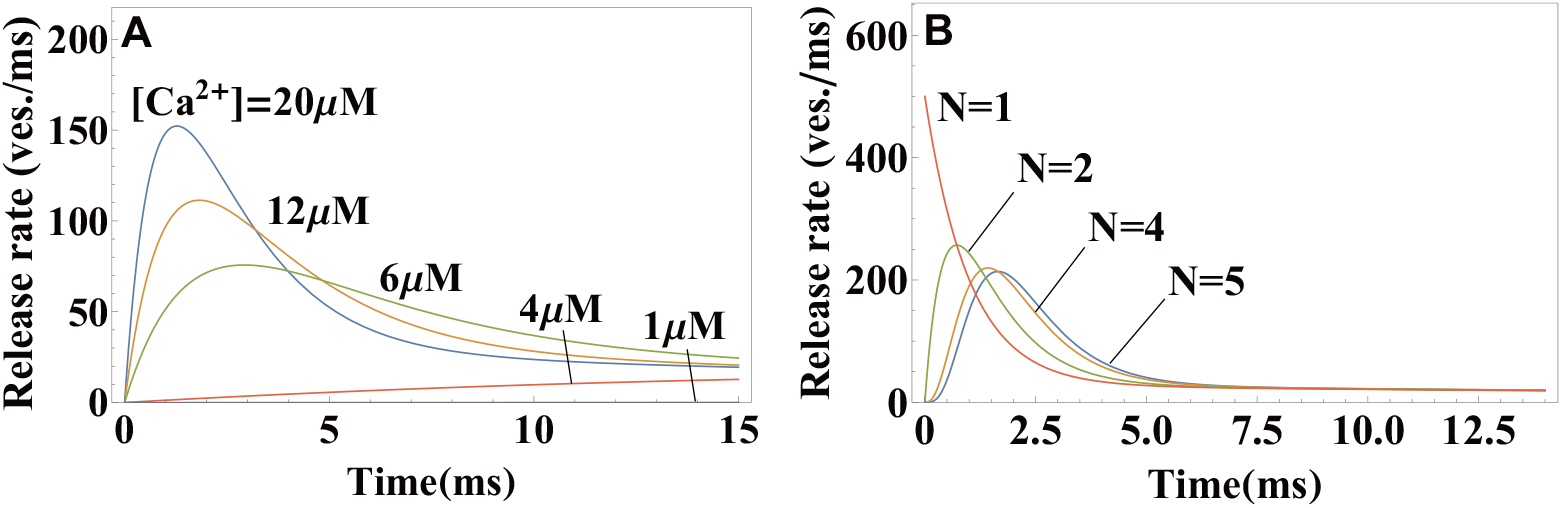
Effects of calcium concentration and the critical number of SNAREs on the neurotransmitter release dynamics, as predicted by the theory. (A) Temporal profiles of the average release rate from Eqs. (2) and (5) in the main text across the range of calcium concentrations (indicated) typical for an action potential. Due to the exponential factor in Eq. (5), the release turns on rapidly upon calcium influx and terminates rapidly with calcium depletion, resulting in a high temporal precision of synaptic release. (B) Temporal profile of the average release rate (Eq. (2)) when *N* SNAREs per vesicle are required for fusion. *N* = 2 provides the optimal balance between stability with respect to fluctuations in calcium concentration (low release rate at sub-millisecond timescale) and temporal precision (the fastest rise of average release rate). Parameter values are given in Appendix 3.

### Universal Scaling Form for the Peak Release Rate

Combining Eq. (4) and Eq. (39), we have

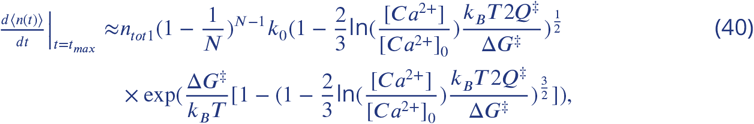

where 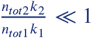 allowed us to ignore the second term in Eq. (4).

Defining 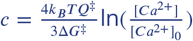 and 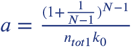, the above equation becomes

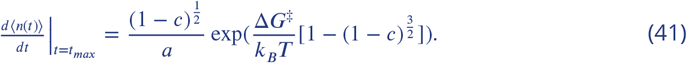

Let 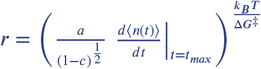. Equation (41) then gives

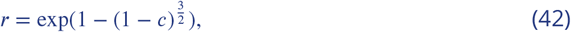

which is Eq. (7) in the main text.

### Peak postsynaptic current and cumulative release

Following the conductance-based model for postsynaptic response (***Destexhe et al., 1994***), the postsynaptic current caused by an ion channel of a given type can be written as

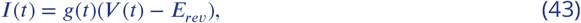

where *g*(*t*) is the conductance of the channel, *V* (*t*) is the postsynaptic membrane potential and *E*_*rev*_ is the reversal potential of the ion corresponding to the ion channel. Different types of channels may have different *g*(*t*) and *E*_*rev*_. The peak value of postsynaptic current is usually dominated by a single type of channel, e.g. AMPA receptor for excitatory synapses or GABA receptor for inhibitory synapses. Thus, in the following, the postsynaptic current will be assumed to be caused by the dominate channel type.

For an action potential triggered at *t* = *t*_*s*_, the conductance has a pulse at *t*_*s*_ of amplitude proportional to the number *n*(*T*) of neurotransmitters released during the action potential:

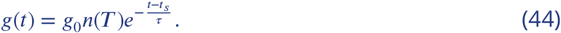

Here, *g*_0_ depends on the intrinsic properties of the ion channel and the channel density, and *τ* is the relaxation timescale of the channel. Since we are concerned with the response to a few (probably one or two) action potentials, Eq. (44) assumes that the postsynaptic receptors are not saturated. Since the time scale for an action potential *T* ~ 1*ms* is much shorter than the relaxation time scale of the ion channel (*τ*_*AMPA*_, *τ*_*GABA*_ ~ 20*ms* − 30*ms* (***Destexhe et al., 1994***)), the action potential can be regarded as a delta-function pulse.

The membrane voltage is usually far from the reversal potential when responding to a few action potentials. According to Eq. (43), the current *I*(*t*) is therefore proportional to the conductance *g*(*t*), and the peak of *g*(*t*) is at *t* = *t*_*s*_. The peak value of postsynaptic current is

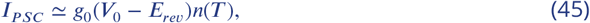

where *V*_0_ is the resting membrane potential. By letting *γ* = *g*_0_(*V*_0_−*E*_*rev*_) and taking the average of both sides of the above equation, we obtain Eq. (8) in the main text.

### Calcium and RRP vesicle kinetics in short-term plasticity

Calcium concentration is assumed to follow the first-order kinetics with the relaxation timescale *τ*_*Ca*_ and a spike-triggered flux:

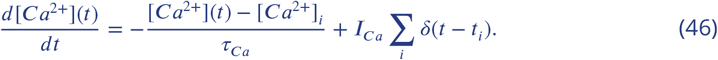

Here, *t*_*i*_ is the presynaptic spike time, and the initial calcium concentration is assumed to be the resting value [*Ca*^2+^]_0_. Calcium concentration immediately after the first spike is then [*Ca*^2+^]_*i*_ = [*Ca*^2+^]_0_ + *I*_*Ca*_.

The number of RRP vesicles is assumed to follow the first-order replenishment kinetics:

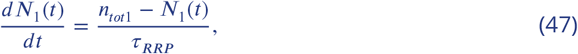

where the RRP pool of total capacity *n*_*tot*1_ is assumed to be full initially: *n*_1,*i*_ ≡ *N*_1_(0) = *n*_*tot*1_.

For a pair of spikes with interpulse interval *τ*_*int*_, the above equations yield solutions for the number of RRP vesicles and calcium concentration immediately after the second spike:

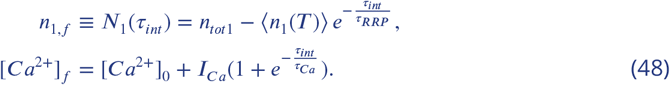

These solutions are used to obtain Eq. (9) in the main text.

The *Ca*^2+^-sensitivity of a SNARE is defined as the ratio of the conformational rate constants (Eq. (39)) during the action potential and at rest: 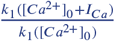.

**Appendix 1 Figure 2.**
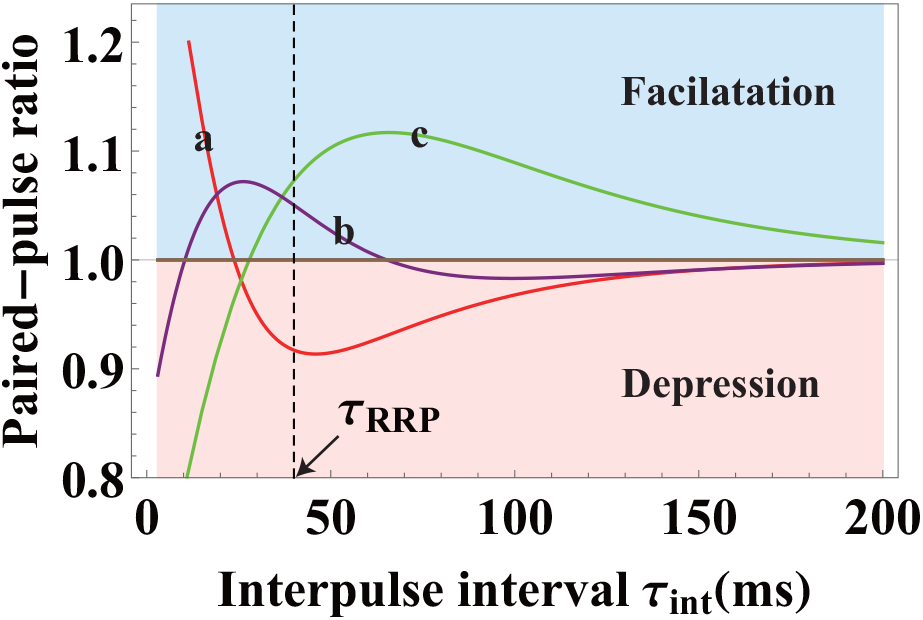
Distinct short-term facilitation/depression modes in synapses that differ on the molecular level, from theory (Eq. (9)). Three different sets of parameters {Δ*G*^‡^, *Q*^‡^, *k*_0_} and *τ*_*Ca*_ are used for curves a, b and c, representing different properties of the molecular constituents for the three synapses. In curve a, the high frequency transient input (with small *τ*_*int*_) is facilitated and the low frequency input (with large *τ*_*int*_) is depressed. The effects are reversed in curve c with depression at high frequency and facilitation at low frequency. In curve b, inputs with intermediate frequencies are facilitated and inputs with high and low frequencies are depressed. The facilitation and depression effects shown here can be amplified if different synapses have different timescales for RRP replenishment (*τ*_*RRP*_). Parameter values are given in Appendix 3.

### Optimal Synaptic Strength

We derive the condition for optimal RRP size as follows. Typically *k*_2_*T* ≪ 1, hence we can ignore the contribution of the reserve pool. The probability that there is no response in the postsynaptic neuron for an action potential of duration *T* can be written as

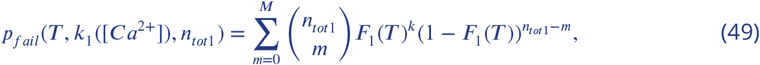

where [*Ca*^2+^] is calcium concentration in pre-synaptic neuron and *F*_1_(*T*) is defined in Eq. (16).

The error probability can be written as

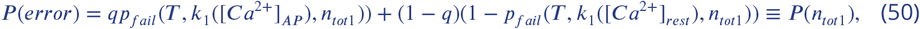

as shown in Eq. (12) in the main text. Here we focus on the dependence of *P*(*error*) on *n*_*tot*1_.

To see whether *P*(*n*_*tot*1_) has a minimum, we solve the following inequality:

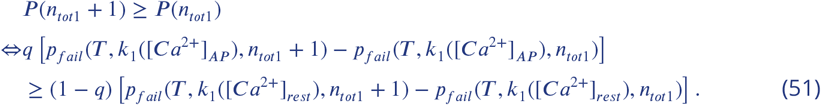

But

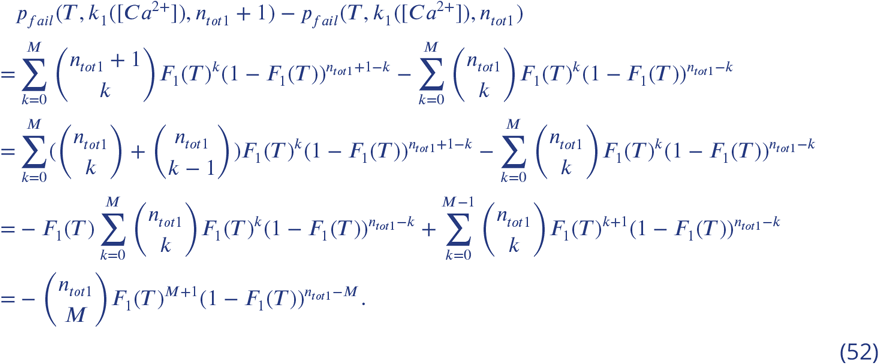

Therefore, Eq. (51) becomes

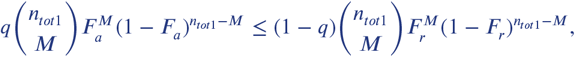

where 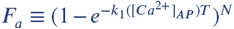 and 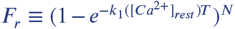 are the probabilities that a vesicle in RRP is fused during the action potential and at rest, respectively.

We can solve *n*_*tot*1_ from the above inequality as follows:

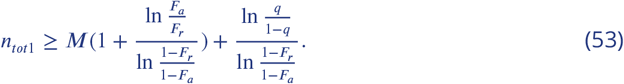

Since *n*_*tot*1_ is an integer, the optimal RRP size 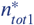 can be written as

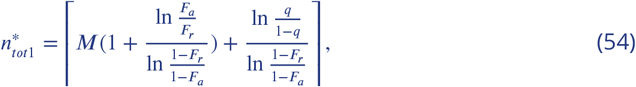

which is Eq. (13) in the main text.

## Appendix 2

### Simulations Set-up

We use Gillespie algorithm to model the fusion dynamics in the simulations. To validate the analytic expressions derived within the framework of the model in Eq. (1) and Fig. 1 in the main text, we performed simulations of this model and examined whether the analytical expressions can accurately recover the input parameters of the simulations when used as a fitting tool. Next, to test the limitations of the assumptions of our model, we performed a modified set of simulations and compared their results with the original model. Specifically, the simulations were modified to incorporate the effects that are thought to be relevant for synaptic transmission *in vivo*: (i) the finite-capacity effect of the readily-releasable pool and (ii) the spatial coupling between voltage-gated calcium channels and vesicle release sites.

### Testing the Analytic Theory

Numerical simulations were carried out for the vesicle fusion model in Eq. (1) and Fig. 1 in the main text with the following parameter values: *n*_*tot*1_ = 500, *n*_*tot*2_ = 1000, *k*_2_ = 0.027*ms*^−1^, Δ*G*^‡^ = 18.4*k*_*B*_*T*, *Q*^‡^ = 1.74, *k*_0_ = 1.67 × 10^−7^*ms*^−1^, and *N* = 2. Calcium concentration was varied from 0.05*μM* to 20*μM*, with the reference value set at [*Ca*^2+^]_0_ = 50*nM*. The number of vesicles that fuse by time *t*, *n*(*t*), was recorded from *t* = 0*ms* to *t* = 100*ms* with time interval 0.4*ms*, and the average over 40 trajectories was calculated to obtain the average cumulative release ⟨*n*(*t*)⟩. The data generated through simulations were then fitted with the analytic expression for the average cumulative release (Eq. (3) in the main text). The theory was found to accurately reproduce the input parameters used in the simulations (Appendix 2 Fig. 1). For low calcium concentration ([*Ca*^2+^] < 1*μM*), the recording time for ⟨*n*(*t*)⟩ had to be extended to ~ 1000*ms* in order to reliably extract the rate constant *k*_1_. This is because, at low calcium concentration, *k*_1_ and *k*_2_ become comparable and the term *k*_1_ − *k*_2_ in the denominator in the expression of *n*(*t*) (Eq. (3) in the main text) tends to cause numerical instability. Nevertheless, the fit to *k*([*Ca*^2+^]) in Eq. (5) in the main text was found to be always reliable as long as there are enough data points at high calcium concentrations ([*Ca*^2+^] > 1*μM*), which is the case for the experimental data in Fig. 2 in the main text.

**Appendix 2 Figure 1.**
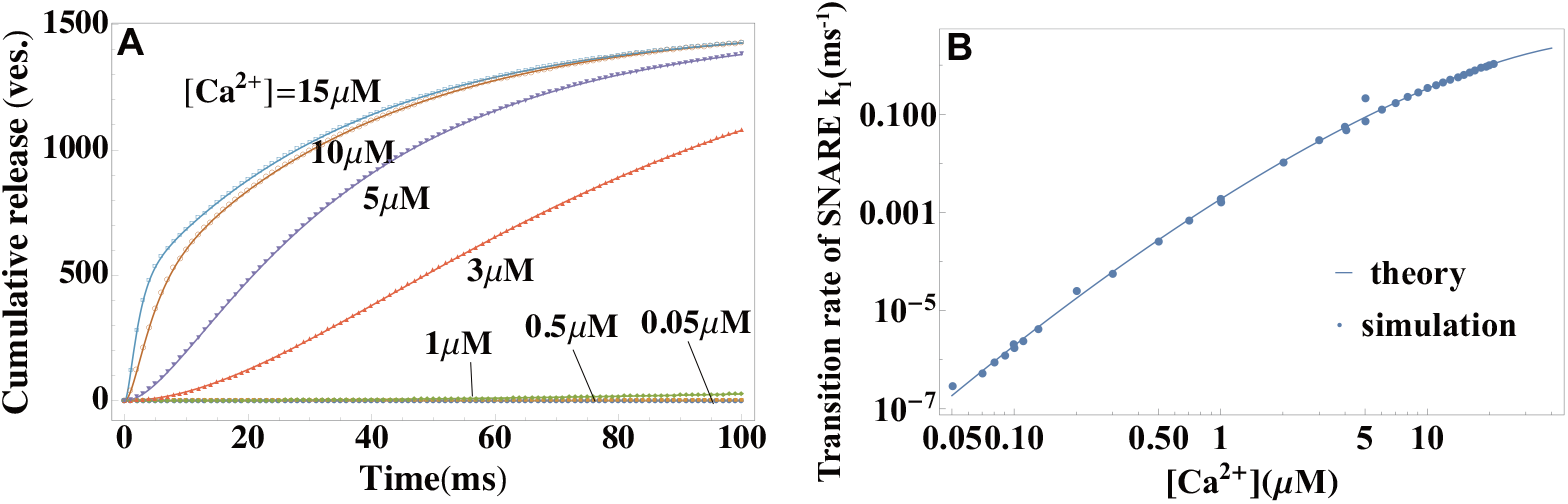
Validation of the theory through simulations. (A) Temporal profiles of average cumulative release at different values of calcium concentrations from simulations (symbols) and a fit to the theory in Eq. (3) in the main text (lines). (B) Calcium dependence of the rate constant of SNARE conformational change from simulations in (A), and a fit to Eq. (5) in the main text. The reference concentration [*Ca*^2+^]_0_ = 50*nM* is set at a typical resting value of a synapse *in vivo* (***Kaeser and Regehr, 2014***) and *N* = 2. The fit yields the height and width of the activation barrier and the rate constant of SNARE conformational change at [*Ca*^2+^]_0_, which accurately recover the input parameters of the simulations. Parameters are listed in Appendix 3.

### Finite-capacity effect of the readily releasable pool

As found in recent experiments (***Biederer et al., 2017***), synaptic vesicles are released at specialized sites, known as active zones, at the presynaptic terminal. Because there are only a finite number of active zones in each presynaptic button, the maximal number of docked vesicles (state *D* in Fig. 1D) is finite. Let *n*_*max*_ be the number of release sites on the presynapic membrane and set *n*_*tot*1_ = *n*_*max*_ in the simulations, which corresponds to the release sites being initially fully occupied by the docked vesicles. To incorporate this finite-capacity effect in the simulations, we now assume that the vesicles in the reserve pool (state *R* in Fig. 1D) can be docked to presynaptic membrane only if there is a vacant release site (*n*_*max*_ − *n*_1_(*t*) > 0).

We note that the model with the finite size of the readily releasable pool corresponds to the *G*/*G*/*N*/*N*/*k* queue model in queueing theory (***Gautam, 2012***). Few results are known for the general *G*/*G*/*N*/*N*/*k* queue model, although bounds and approximation methods have been developed for various situations. When the capacity *N* → +∞, the *G*/*G*/*N*/*N*/*k* queue model converges to our model in Appendix 1 with infinite capacity of the RRP pool, and is exactly solvable. Here, rather than seeking analytic approximations for the finite-capacity effect, we use simulation to explore its properties and the validity of our model in the light of this effect. We define the dimensionless ratio 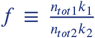 and change it from 10 to 0.5. When *f* ≫ 1, the depletion rate of the readily releasable pool is much larger than the replenishing rate, and the readily releasable pool is effectively of an infinite capacity. As expected, the theory (Eq. (3) in the main text) is valid in this regime (Appendix 2 Fig. 2A). When *f* decreases, deviations from the analytic theory appear, with the cumulative release being slower than that predicted by Eq. (3) in the main text (Appendix 2 Fig. 2A). In real neuron, action potential-evoked calcium elevation will lead to *k*_1_ ≫ *k*_2_ and thus *f* ≫ 1, therefore, Eq. (3) in the main text is expected to perform well in the biologically relevant range of parameters.

**Appendix 2 Figure 2.**
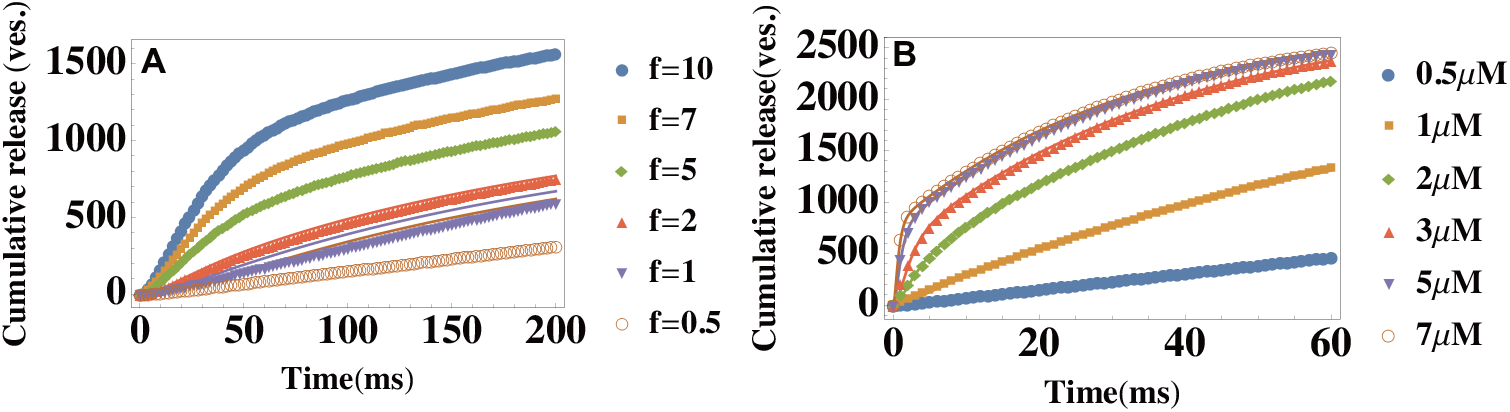
Testing the limitations of the theory: Finite capacity of the readily releasable pool and heterogeneity in vesicle pools. (A) Temporal profiles of the average cumulative release *n*(*t*) at different ratios 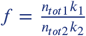. Parameter values are *n*_*tot*2_ = 1000, *k*_2_ = 0.005*ms*^−1^ and *k*_1_ = 0.05*ms*^−1^, and *n*_*tot*1_ is varied to change the ratio *f* of the depletion and replenishment rates of the readily releasable pool. Symbols: data generated from modified simulations that introduced the finite-capacity effect of the readily releasable pool, lines: Eq. (3) in the main text with the same parameter values as those used in the simulations. (B) Temporal profiles of the average cumulative release *n*(*t*) at different values of the extracellular calcium concentration [*Ca*^2+^]_*out*_. The parameters are described in the text. [*Ca*^2+^]_*out*_ is shown in the legend. Data is fitted with Eq. (3) in the main text. The cumulative release rate exhibits a double-exponential shape if *λ*_*max*_ − *λ*_*min*_ is not too large (≲ 10*nm*).

### The Effect of Heterogeneity among Release Sites

It has been pointed out that the docked vesicles in the same synapse may have different release rates due to their different distances to the voltage-gated calcium channels (***Trommershäuser et al., 2003; Neher, 2015***). Action potential-evoked calcium influx forms a so-called nanodomain around each channel. Let us assume that diffusion and buffering are the dominant factors that shape the concentration profile of calcium. When the channel is open, the steady state of calcium concentration profile can be described by the following reaction-diffusion equation:

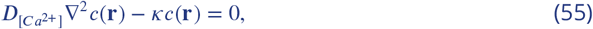

where 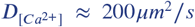 is the diffusion coefficient of a calcium ion inside the cell and *κ* = *k*_[*B*] ≈ 0.3*μs*^−1^ (***Delvendahl et al., 2015***) is the binding rate that characterizes calcium buffers. Eq. (55) can be solved by assuming spherical symmetry (***Neher, 1998***):

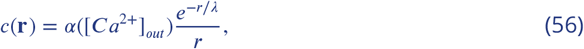

where 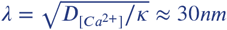 sets the characteristic length scale of the calcium nanodomain, *α*([*Ca*^2+^]_*out*_) measures the magnitude of the calcium current through the channel at extracellular calcium concentrations [*Ca*^2+^]_*out*_, and *r* is the distance from the channel. Due to the decaying concentration profile in Eq. (56), vesicles that are closer to the channel experience higher calcium concentration and thus have higher release rate. The relative positions of the release site and voltage-gated calcium channels on the presynaptic membrane may therefore have a significant impact on the action-potential-evoked vesicle release dynamics.

Recently, several studies established the nanoscale organization of the molecular apparatus around the vesicle release sites (***Stanley, 2016; Gramlich and Klyachko, 2019; Biederer et al., 2017***). It has been found that release sites and channels together form clusters within active zones on the presynaptic membrane (***Maschi and Klyachko, 2017; Miki et al., 2017; Nakamura et al., 2015***). The typical size of an active zone is about 250nm (***Gramlich and Klyachko, 2019***), with multiple release sites present within a single active zone (***Maschi and Klyachko, 2017***). Multiple channels cluster around a single release site and their distances to the release site are regulated by scaffold proteins (***Böhme et al., 2016***).

Based on these experimental facts, we set up our modified simulations as follows. Each active zone is modeled as a disk of radius *r* = 125nm, the total number of active zones on the presynaptic synapse is *N*_*AZ*_ = 300, the number of release sites in an active zone is *N*_*r*_ = 3 and the number of channels around each release site is *N*_*c*_ = 3. These values are chosen to mimic the organization of release sites in the Calyx of Held (***Borst and Soria van Hoeve, 2012***). We further assume that the release sites are uniformly distributed within each active zone (***Maschi and Klyachko, 2017***), and the channels are uniformly distributed around each release site within the range of distances (*λ*_*min*_ = 30*nm, λ*_*max*_ = 40*nm*). The calcium current 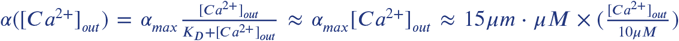 (***Schneggenburger et al., 1999***), and the concentration [*Ca*^2+^]_*out*_ is varied from 0.5*μM* to 10*μM*. The release rates for docked vesicles are determined by Eq. (5) in the main text and the parameter values are the same as in the above simulations (See “Testing the Analytic Theory”). Each release site is assumed to have its corresponding reserve pool of size *n*_*tot*2_ = 3, and the release rate for vesicles in the reserve pool is *k*_2_ = 0.02*ms*^−1^. Simulation results and the corresponding fits to Eq. (3) in the main text are shown in Appendix 2 Figure 2B. The simulations show that, as long as *λ*_*max*_ − *λ*_*min*_ is not too large, our theory is accurate. Further simulations (not shown) show that, if *λ*_*max*_ − *λ*_*min*_ is too large, the cumulative release curve in Appendix 2 Figure 2B is no longer double-exponential. The fact that cumulative release has in fact been observed to be double-exponential (***Miki et al., 2018***) indicates that the distance between the channel and the release sites is likely to be tightly regulated by the scaffold proteins. We conclude that, in the range of parameters that correspond to real biological systems, the spatial heterogeneity of calcium concentration has no significant effect on the accuracy of the results presented in the main text.

## Appendix 3

### Critical Number of SNAREs, *N*

To achieve robustness of the fits with limited experimental data available, the fits in the main text were performed at a fixed value of the critical number of SNAREs, *N*, necessary for vesicle fusion. The choice of *N* = 2 was based on the indirect experimental evidence (***Sinha et al., 2011***). Additionally, we performed the least square fits of the *in vivo* data in Fig. 2 in the main text with Eq. (3) with the values of *N* = 1, 2, 3, 4, 5. Values *N* ≥ 6 would be too large to be biologically realistic (***Brunger et al., 2018***). The fitting errors for different values of *N* are reported in Appendix 3 Table 1.

It can be seen that *N* = 2 results in the consistently smallest fitting errors across all values of calcium concentrations used in the experiment, and thus represents the optimal fit. This numerical test provides an additional support for *N* = 2 representing the critical number of SNAREs necessary for fusion.

**Appendix 3 Table 1.** Errors from the least square fit for different values of *N*. The fits are performed for the experimental data from (***Kochubey et al., 2009***) to Eq. (3) in the main text. The fitting error is calculated according to **Σ**_*i*_ |*y*_*i*_ − *f*(*x*_*i*_)|^2^. The fit with *N* = 2 results in consistently smallest fitting errors across all calcium concentrations used in the experiment.

| Least Squares Error | $N = 1$ | $N = 2$ | $N = 3$ | $N = 4$ | $N = 5$ |
| --- | --- | --- | --- | --- | --- |
| $[Ca^{2+}] = 27\mu M$ | 27.431 | 25.896 | 26.742 | 29.016 | 33.234 |
| $[Ca^{2+}] = 11\mu M$ | 19.662 | 13.968 | 14.656 | 15.374 | 16.737 |
| $[Ca^{2+}] = 7\mu M$ | 10.241 | 8.115 | 8.189 | 8.260 | 8.416 |
| $[Ca^{2+}] = 6\mu M$ | 84.908 | 84.732 | 85.255 | 85.553 | 85.701 |

### Parameter Values Extracted from the Fits or Used for Illustration

**Appendix 3 Table 2.** Microscopic parameters of synaptic fusion machinery extracted from the fits in Fig. 2A of the main text.

| $\Delta G^\ddagger (k_B T)$ | $Q^\ddagger$ | $\log_{10}(\frac{n_{tot} k_0}{ms^{-1}})$ | Data Source |
| --- | --- | --- | --- |
| $34.3 \pm 3.0$ | $4.04 \pm 0.48$ | $-10.8 \pm 1.3$ | Calyx of Held (Schneggenburger and Neher, 2000) |
| $35 \pm 19$ | $2.24 \pm 2.94$ | $-6.6 \pm 0.84$ | Calyx of Held (Lou et al., 2005) |
| $27.0 \pm 1.31$ | $3.11 \pm 0.25$ | $-10.1 \pm 0.5$ | Calyx of Held (Bollmann et al., 2000) |
| 16.6 | 1.33 | -3 | PF-MLI (Miki et al., 2018) |
| $7.63 \pm 3.34$ | $1.67 \pm 1.18$ | $-1.33 \pm 0.66$ | Photoreceptor (Duncan et al., 2010) |
| $19.8 \pm 1.5$ | $1.77 \pm 0.5$ | $-4 \pm 1.2$ | Calyx of Held (Wölfel et al., 2007) |
| $20.3 \pm 1.5$ | $1.93 \pm 0.2$ | $-4.6 \pm 0.6$ | Calyx of Held P89 (Kochubey et al., 2009) |
| $25.2 \pm 1.9$ | $2.64 \pm 0.27$ | $-6.5 \pm 0.7$ | Calyx of Held P12 (Kochubey et al., 2009) |
| 20 | 1.41 | -8.6 | *in vitro* (Diao et al., 2012) |

In Fig. 2C, the fit yields the height and width of the activation barrier and the rate constant of the SNARE conformational change in the resting state ([*Ca*^2+^]_0_ = 50*nM*): Δ*G*^‡^ (18 ± 1.0)*k_B_T*, *Q*^‡^ 1.77 ± 0.13 and 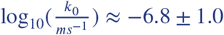.

In Fig. 2D, the fit yields *k*_1_ ≈ 2.32*s*^−1^ and *k*_*2*_ ≈ 0.0117*s*^−1^ (250*μ*M) and *k*_1_ ≈ 2.17*s*^−1^ and *k*_2_ ≈ 0.0198*s*^−1^ (500*μM*).

In Fig. 2E, parameter values used: Δ*G*^‡^ = 35*k*_*B*_*T*, *Q*^‡^ = 1, *k*_0_ = 10^−5^*s*^−1^, 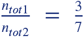 and *k*_2_ = 0.0105*ms*^−1^.

In Fig. 3A, parameter values used: *T* = 3*ms*, Δ*G*^‡^ = 18.5*k*_*B*_*T*, *Q*^‡^ = 1.77, *k*_0_ = 1.67 × 10^−7^*ms*^−1^, *n*_*tot*1_ = 1000, *τ*_*RRP*_ = 40*ms*, [*Ca*^2+^]_0_ = 50*nM* and *I*_*Ca*_ = 10*μM*.

In Fig. 3B, the values for *T*, *Q*^‡^, *n*_*tot*1_, *τ*_*RRP*_, [*Ca*^2+^]_0_ and *I*_*Ca*_ are the same as Fig. 3A. Δ*G*^‡^ = 15.4*k*_*B*_*T* and *k*_0_ = 5.45 × 10^−7^*ms*^−1^ are changed such that the *Ca*^2+^-sensitivity becomes 1/3 while the response to an single action potential is the same as Fig. reffig:shortermA.

In Fig. 3C, parameter values used: *k*_1_ = 0.5*ms*, *M* = 100.

In Fig. 3D, parameter values used: *M* = 10, *k*_1_([*Ca*^2+^]_*a*_) = 0.32*ms*^−1^, *q* = 10^−1^, *T* = 2.5*ms*.

In Appendix 1 Figure 1A, parameter values are *k*_0_ = 2.3 × 10^−3^*ms*^−1^, Δ*G*^‡^ = 14.77*k*_*B*_*T*, *Q*^‡^ = 3.34, *n*_*tot*1_ = 1000, *n*_*tot*2_ = 1000, *k*_2_ = 0.027*ms*^−1^, *N* = 2. Parameters values used in Figure 1B is the same as Figure 1A except *k*_1_ = 1*ms*^−1^.

In Appendix 1 Figure 2, parameter values are Δ*G*^‡^ = 18.5*k*_*B*_*T*, *Q*^‡^ = 1.77, *k*_0_ = 1.67 × 10^−7^*ms*^−1^ and *τ*_*Ca*_ = 12*ms* for curve a; Δ*G*^‡^ = 16.5*k*_*B*_*T*, *Q*^‡^ = 1.57, *k*_0_ = 1.4 × 10^−6^*ms*^−1^ and *τ*_*Ca*_ = 48*ms* for curve b; Δ*G*^‡^ = 20*k*_*B*_*T*, *Q*^‡^ = 2, *k*_0_ = 4 × 10^−8^*ms*^−1^ and *τ*_*Ca*_ = 48*ms* for curve c. For all three curves, *T* = 3*ms*, *n*_*tot*1_ = 1000, *τ*_*RRP*_ = 40*ms*, [*Ca*^2+^]_0_ = 50*nM* and *I*_*Ca*_ = 10*μM*.

In Appendix 2 Figure 1, the fit yields the height and width of the activation barrier and the rate constant for the SNARE conformational change at [*Ca*^2+^]_0_: Δ*G*^‡^ = (18.5 ± 0.11)*k*_*B*_*T*, *Q*^‡^ = 1.74 ± 0.01 and *k*_0_ = (1.88 ± 0.07) × 10^−7^*ms*^−1^. The fitting parameters accurately recover the input parameters of the simulations: Δ*G*^‡^ = 18.4*k*_*B*_*T*, *Q*^‡^ = 1.77 and *k*_0_ = 1.67 × 10^−7^*ms*^−1^.

